# RhoA effectors LOK/SLK activate ERM proteins to locally inhibit RhoA and define apical morphology

**DOI:** 10.1101/2020.07.02.185298

**Authors:** Riasat Zaman, Andrew Lombardo, Cécile Sauvanet, Raghuvir Viswanatha, Valerie Awad, Locke Ezra-Ros Bonomo, David McDermitt, Anthony Bretscher

**Author notes:** These authors contributed equally to this work (Riasat Zaman, Andrew Lombardo). Address for correspondence (A.B.).

## Abstract

Activated Ezrin-Radixin-Moesin (ERM) proteins link the plasma membrane to the actin cytoskeleton to generate apical structures, including microvilli. Among many kinases implicated in ERM activation are the homologs LOK and SLK. CRISPR/Cas9 was used to knockout all ERM proteins or LOK/SLK in human cells. LOK/SLK knockout eliminates all ERM activating phosphorylation. The apical domain of cells lacking LOK/SLK or ERMs is strikingly similar and selectively altered, with loss of microvill, and junctional actin replaced by ectopic myosin-II containing apical stress-fiber-like structures. Constitutively active ezrin can reverse the phenotypes of either ERMs or LOK/SLK knockouts, showing that the major function of LOK/SLK is to activate ERMs. Both knockout lines have elevated active RhoA with concomitant enhanced myosin light chain phosphorylation, revealing that active ERMs are negative regulators of RhoA. As RhoA-GTP activates LOK/SLK to activate ERM proteins, the ability of active ERMs to negatively regulate RhoA-GTP represents a novel local feedback loop necessary for the proper apical morphology of epithelial cells.

## Introduction

Essentially all eukaryotic cells are polarized, or have the potential to become polarized. This requires local regulation to define the morphology and composition of each cellular domain. A particularly well studied example is the intestinal epithelial cell with a highly ordered apical domain displaying abundant microvilli and having a protein and lipid composition distinct from the more planar basolateral membrane. While much is known about extrinsic cues that instruct the cell to polarize (Rodriguez-Boulan and Macara, 2014), much less is known about how the morphology of the individual domains is regulated. To address this, we have studied how the apical domain of epithelial cells is regulated to assemble bundles of actin filaments that support the plasma membrane of microvilli, in contrast to the flatter structure of the basolateral membrane.

The structural integrity of apical microvilli requires active ezrin/radixin/moesin (ERM) proteins (Fehon et al., 2010). These proteins exist in a cytoplasmic closed state and an active open conformation where the N-terminal FERM domain binds the plasma membrane and the C-terminal F-actin binding domain binds the underlying actin filaments (Gary and Bretscher, 1995). Activation requires phosphorylation of a conserved threonine, T567 in ezrin (Hayashi et al., 1999; Matsui et al., 1998). The physiologically relevant kinase(s) responsible for ERM phosphorylation have been investigated for decades. Among the many kinases suggested are Rho kinase (Haas et al., 2007; Matsui et al., 1998; Tran Quang, 2000), PKCα (Ng et al., 2001); PKCθ (Pietromonaco et al., 1998), MST4 (Gloerich et al., 2012; ten Klooster et al., 2009), NcK-interacting kinase (Baumgartner et al., 2006) and LOK (Belkina et al., 2009; Viswanatha et al., 2012). More recent evidence has suggested that the related LOK and SLK are significant vertebrate kinases for ERM phosphorylation (Viswanatha et al., 2012), being the homologs of Slik that is responsible for phosphorylating the single ERM in flies, moesin (Hipfner et al., 2004). LOK and SLK belong to the germinal center-like kinase (GCK)-V subfamily of kinases (Kuramochi et al., 1997). They consist of a conserved N-terminal kinase domain, a less-conserved intermediate region and a moderately conserved C-terminal domain. The LOK C-terminal domain inhibits the kinase activity of LOK in cells, most likely through a cis-interaction, as well as targeting LOK to the apical membrane (Pelaseyed et al., 2017; Viswanatha et al., 2012). To maintain a strictly apical distribution, ezrin has to undergo phosphorylation/dephosphorylation cycles in which it is locally phosphorylated and subject to dephosphorylation by delocalized phosphatase Mypt1/PP1. As a result, constitutively active ezrin is unable to maintain apical restriction (Viswanatha et al., 2012).

The Rho family of small GTP-binding proteins are major regulators of microfilaments (Hall and Nobes, 2000). In early work Speck et al. (Speck et al., 2003) found that defects in fly moesin could be counteracted by antagonizing Rho activity, suggesting that ERM proteins might be able to regulate contractility in some manner. RhoA is capable of binding to a diverse range of effectors that influence the reorganization of actin into stress fibers and focal adhesions (Hall, 1998). Among these, the effector ROCK modulates microfilament organization and function in at least two distinct ways. First, it activates myosin II by directly phosphorylating the myosin regulatory light chain (Kimura et al., 1996) and inactivating myosin light chain phosphatase (Mypt1/PP1) (Kimura et al., 1996). Second, ROCK stabilizes F-actin by phosphorylating LIM kinase to abrogate its inhibitory activity towards cofilin, an F-actin destabilizing factor (Arber et al., 1998; Maekawa et al., 2017; Yang et al., 1998). The net effect results in an increase of cortical tension that can drive cellular contraction. ERM proteins have long been suggested to be downstream targets of ROCK, first as direct targets and then as indirect targets through Rho activation of PI-4-P5K (Matsui et al., 1999).

Until very recently, the potential relationship between Rho and LOK and SLK had not been studied. A BioID screen for regulators and effectors of RhoA in cultured cells identified SLK (and LOK) as effectors of RhoA (Bagci et al., 2020). RhoA-GTP was shown to bind to the C-terminal domain of SLK, the corresponding domain that negatively regulates the kinase activity and targets LOK to the apical domain. Moreover, active RhoA can activate SLK in its ability to phosphorylate ezrin. Thus, a direct pathway exists from RhoA-GTP to its effector LOK/SLK to ERM protein activation.

Further uncovering the functions and exploring signaling pathways related to ERM proteins and LOK and SLK in cultured cells has been hampered by redundancy, and the possibility that they perform essential functions. Here we show that it is possible to generate cultured cells lacking all ERM proteins, and cells lacking both LOK and SLK. This has allowed us to explore the roles of these proteins, both by characterizing cells lacking them, but also by re-introduction of variants. Surprisingly, cells lacking LOK and SLK have a strikingly similar phenotype to cells lacking ERM proteins. Our results establish that LOK and SLK are the major, and possibly only, kinases that phosphorylate the conserved threonine of ERM proteins in epithelial cells. Further, rescue experiments indicate that ERM proteins are the major substrates of LOK and SLK that regulate cell morphology. Moreover, loss of either LOK/SLK or ERM proteins selectively modifies the apical actin cytoskeleton of epithelial cells by elevating RhoA signaling to redistribute actin from microvilli and junctions to generate ectopic apical stress fibers. We propose that LOK/SLK and ERM proteins function as a module that defines apical morphology of epithelial cells in a process that involves a negative feed-back loop to locally down-regulate Rho activity.

## Results

### Jeg3 cells lacking ERM proteins or LOK/SLK are viable and lack microvilli

In earlier studies we described human cells modified by CRISPR/Cas9 to lack expression of LOK (Pelaseyed et al., 2017), but so far no cultured cells have been described that genetically lack specific ERM proteins. We set out to determine if cells lacking all ERM proteins, or lacking both LOK and SLK, are viable, and if so, what phenotypes were conferred. Our studies were performed in Jeg3 epithelial cells derived from a human choriocarcinoma as these cells exhibit abundant apical microvilli (Pakkanen et al., 1987). Of the ERM proteins, Jeg3 cells express ezrin and radixin, but not moesin (Figure 1A). We first used CRISPR/Cas9 to isolate cells lacking either ezrin or radixin or SLK (Figure 1A). We then generated pairs of double knockout cells, lacking all detectable ezrin and radixin, or LOK and SLK (Figure 1A). Importantly, moesin was not expressed in the Jeg3 ezrin^-/-^ radixin^-/-^ cells—which can be identified as moesin is expressed in HeLa cells (Figure 1A, lane 1). Thus, the Jeg3 cells lacking all ERM proteins are viable. Likewise, Jeg3 cells lacking both LOK and SLK are viable. However, both ezrin^-/-^ radixin^-/-^ and LOK^-/-^ SLK^-/-^ cells grew slower than their wildtype counterparts (Figure S1A).

**Figure 1:**
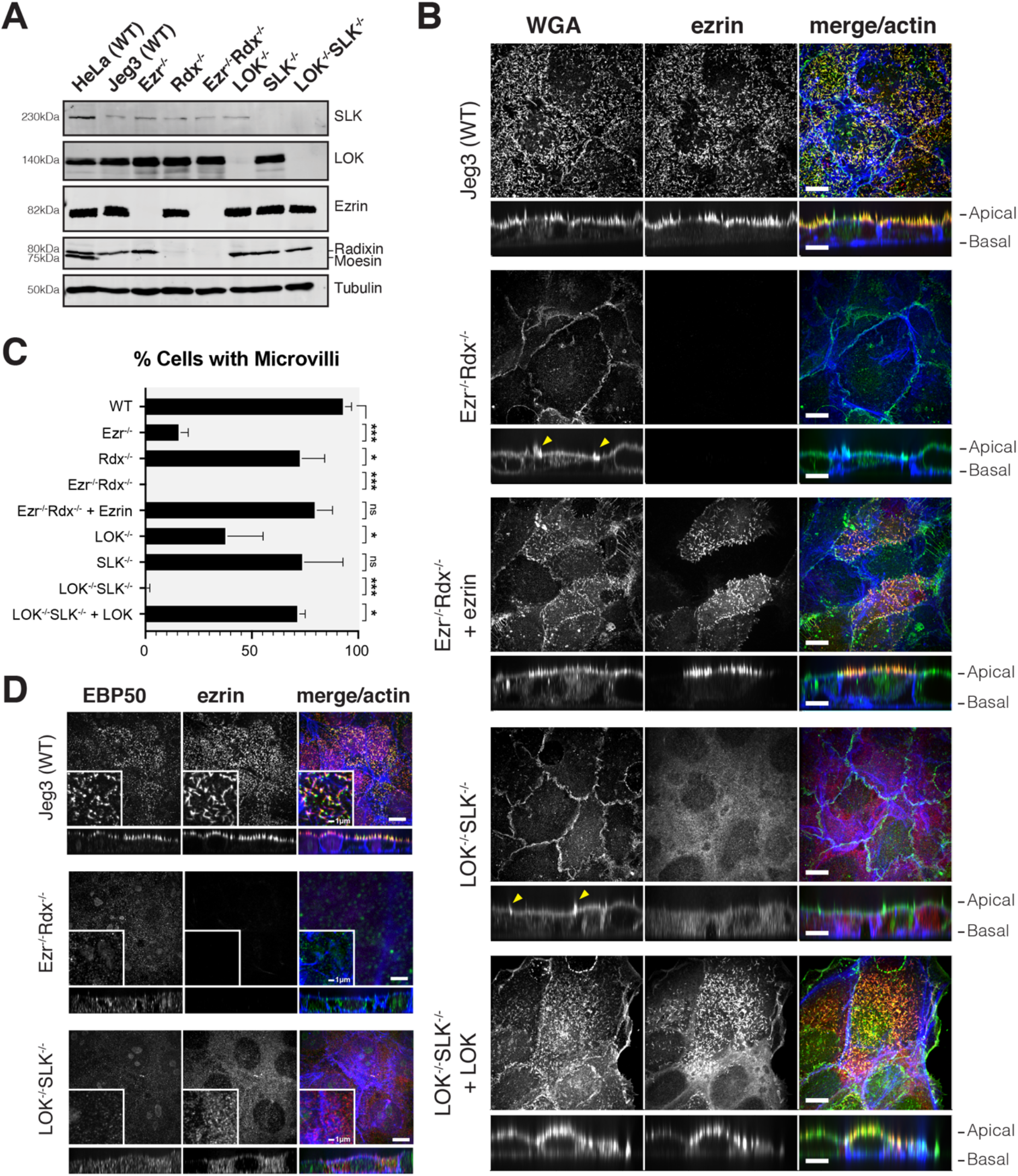
Jeg3 cells lacking ERM proteins or LOK/SLK are viable and lack microvilli. (A) Protein expression using indicated antibodies in western blots of ERM proteins and LOK and SLK in Jeg3 CRISPR cell lysates. HeLa cell lysate was used as a control for expression of moesin, which is not present in Jeg3. Tubulin expression was used as a loading control. (B) Jeg3 cell lines lacking ERM expression or LOK and SLK expression were stained with Wheat-Germ-Agglutinin (WGA), ezrin and actin. Yellow arrows indicate strong membrane ruffles. (C) Percentage of cells expressing microvilli. Bars represents mean±SEM, N=3. Non-significant (ns) p-values are as follows: Ezr ^-/-^ Rdx ^-/-^ + ezrin = 0.0770 ; LOK ^-/-^ SLK ^-/-^ + LOK = 0.2079. (D) Staining of cells with EBP50, ezrin and actin in the indicated cells. Scale bars, 10μm unless otherwise noted. Vertical sections were expanded three-fold for clarity. P-values were calculated with Welch’s t-test (* p≤0.05, ** p≤0.01, *** p≤0.001).

As ERM proteins and LOK and SLK have been implicated in the formation of microvilli on these cells, we examined whether single and double knockout cells retained microvilli. To assess the presence of microvilli, we could not use the traditional ERM proteins as markers, so we utilized labelled wheat germ agglutinin (WGA) that binds to plasma membrane glycoproteins and allows for the identification of cell surface structures. In wildtype cells, WGA colocalizes with ezrin in surface microvilli (Figure 1B). Consistent with earlier reports, individual reduction of ezrin or genetic loss of LOK resulted in a reduction in the number of cells with apical microvilli, whereas loss of radixin or SLK had little phenotype (Figure 1C, Figure S1B). Strikingly ezrin^-/-^ radixin^-/-^ and LOK^-/-^ SLK^-/-^ cells totally lack microvilli, with WGA staining mostly associated with membrane ruffles that form above membrane contact sites (Figure 1B, arrows and 1C). Ezrin is cytosolic in LOK^-/-^ SLK^-/-^ cells, consistent with phosphorylation by LOK/SLK being required to activate it. Stable expression of ezrin in ezrin^-/-^ radixin^-/-^ cells and LOK in LOK^-/-^/SLK^-/-^ restored the presence of microvilli to the cell surface. We conclude that both an ERM protein and LOK or SLK are necessary for the presence of apical microvilli. We also examined the localization of the interactor of active ezrin, EBP50 (ERM-binding phosphoprotein of 50kD)/NHERF1 (Figure 1D). Whereas EBP50 was present in microvilli in wildtype cells, in both double knockout cell lines EBP50 was cytosolic. Both the WGA and EBP50 staining supports the conclusion that the loss of ERMs or LOK/SLK disrupts the presence of microvilli from the apical surface.

### LOK/SLK are the major kinases for ERM phosphorylation

ERM proteins undergo activation by C-terminal phosphorylation to exhibit their membrane-cytoskeletal linking function (Pearson et al., 2000; Turunen et al., 1994). The identity of the relevant kinase has been controversial, so we assessed the contribution of LOK and SLK to ERM phosphorylation in LOK^-/-^ SLK^-/-^ cells. Using an antibody that detects the relevant epitope on all ERM proteins (phosphorylation of T567 in ezrin, T564 in radixin, and T558 in moesin) we found that LOK^-/-^ SLK^-/-^ cells appeared to be entirely devoid of all ERM phosphorylation (Figure 2A). However, because the pERM antibody produces a small background staining to unphosphorylated ezrin (Pelaseyed et al., 2017), we repeated the experiment employing phos-Tag gels in which phosphorylated ERMs migrate slower than their unphosphorylated counterparts. Again, we were unable to detect any phosphorylated ERM proteins in the absence of LOK and SLK (Figure 2B), so LOK and SLK appear to be the only significant ERM kinases in Jeg3 cells.

**Figure 2:**
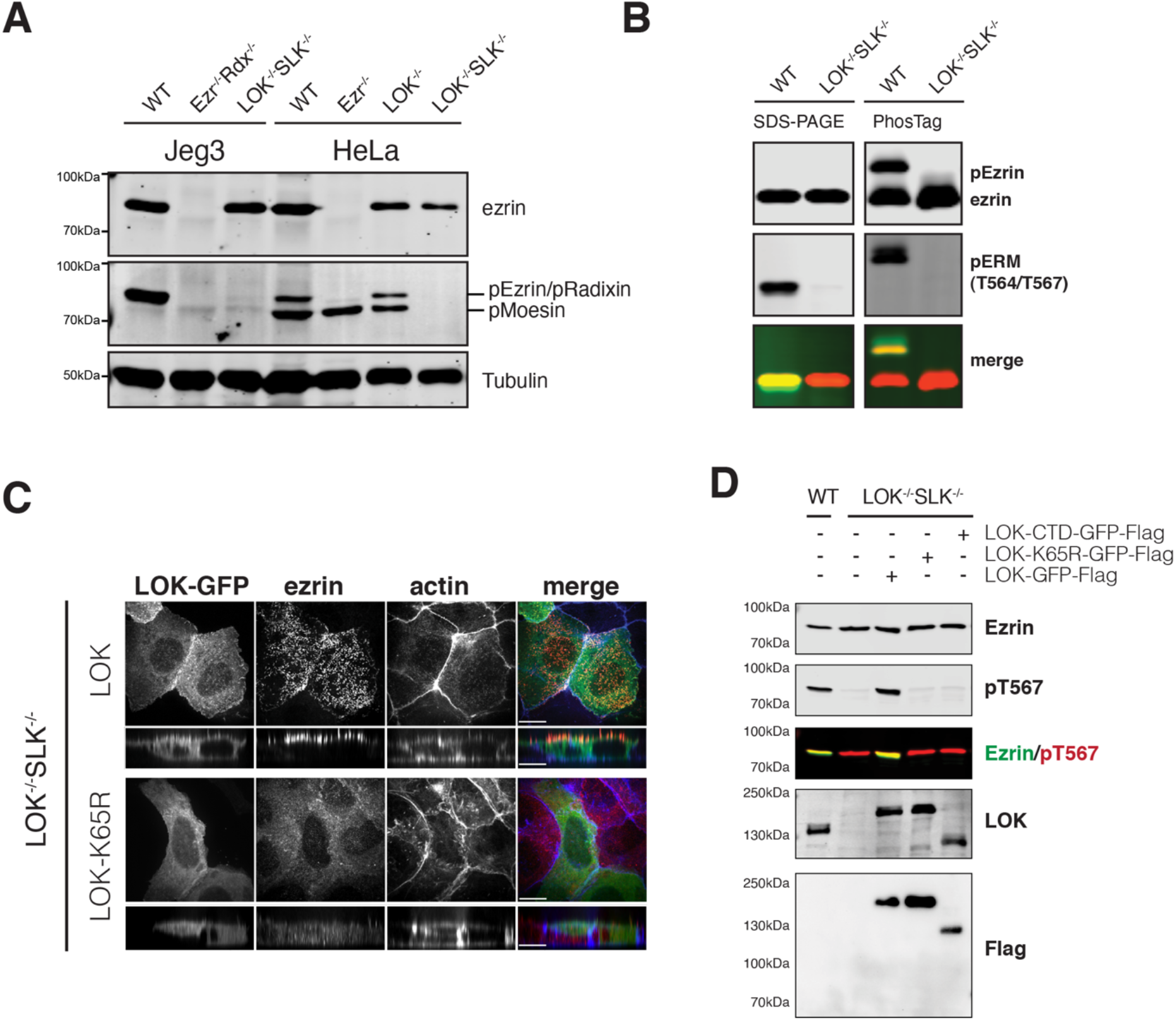
LOK/SLK are the major ERM kinases. (A) Ezrin and phospho-ERM levels in Jeg3 or HeLa knockout cells. Tubulin is used as a loading control. (B) Extracts of wildtype or LOK^-/-^SLK^-/-^ cells were resolved by either 6% SDS-PAGE gel or 6% Mn-Phos-Tag SDS-PAGE gel and blotted for ezrin and phosphoERM. Approximately half of endogenous ezrin is phosphorylated in wildtype cells as seen by the shift in band size. No phospho-shift is detected in LOK^-/-^SLK^-/-^ cells. (C) LOK^-/-^SLK^-/-^ cells transfected with either LOK-GFP-flag or K65R-LOK-GFP-flag and then co-stained with ezrin and actin. K65R-LOK is unable to rescue apical ezrin localization. (D) Extracts of cells transfected with wildtype LOK or LOK mutants were collected and blotted for endogenous ezrin or phosphoERM. Lysates were also blotted for flag or LOK to check expression of the constructs relative to wildtype LOK. Scale bars 10μm.

To explore if this is true in other cells, we also generated LOK^-/-^ SLK^-/-^ knockouts in HeLa cells. While these cells survived clonal isolation and lysate collection, they grew very slowly and could not be maintained. Nevertheless, analysis of cell lysates revealed that in HeLa cells as well, LOK and SLK are the major ERM kinases (Figure 2A). Further, we also tried to knock out LOK and SLK in epithelial Caco-2 cells, which proved inviable past clonal selection indicating the importance of these kinases. Likewise, we tried to isolate HeLa and Caco-2 cells lacking ezrin. Although we nursed the growth of enough cells to demonstrate loss of ezrin, they could not be passaged. These results suggest that ERM proteins are more important for viability in HeLa and Caco-2 cells than in Jeg-3 cells. It was fortuitous that we started with Jeg-3 cells as it has allowed us to use the double knock-outs to explore in more detail the phenotypes conferred by loss of these proteins, or reintroduction of variants.

In transfection-based experiments, expression of LOK-GFP-Flag was able to both restore ERM phosphorylation and apical microvilli to LOK^-/-^ SLK^-/-^ cells (Figures 1B&C, 2D). LOK contains an N-terminal kinase domain and C-terminal domain responsible for both regulating and localizing the kinase (Pelaseyed et al., 2017). As expected, expression of a K65R kinase-inactive variant of LOK was unable to restore phosphorylation or microvilli (Figure 2C,D).

### Absence of activated ERM proteins induces the formation of apical actin/myosin II bundles and alters cell-cell junctions of epithelial cells

To explore what additional phenotypes might be associated with loss of ERM proteins and their activating kinases, we examined the structure of cell junctions and the actin distribution in the knockout cells. A characteristic feature of epithelial cells is their ability to form strong cell-to-cell contact sites. Using the tight junction marker ZO-1, we observe a remarkably similar phenotype in ezrin^-/-^ radixin^-/-^ and LOK^-/-^ SLK^-/-^ cells. In both cases, the cells maintain cell-to-cell contacts, but the contact sites are uneven and occasionally have breaks in their ZO-1 and greatly reduced junctional actin staining (Figure 3A, magenta arrows). In contrast, wildtype Jeg3 cells form uniform and relatively linear contacts along cell junctions co-staining with actin (Figure 3A, blue arrows). By determining the ratio between the contour length and shortest distance in ZO-1 staining from a three-cell junction to the next, we quantified the straightness of the tight junctions in cells under various conditions (Figure 3B). This measurement of tortuosity was slightly above 1.0 for wildtype cells, reflecting their almost linear nature. For either ezrin^-/-^ or LOK^-/-^ this rose to 1.15, and for ezrin^-/-^ radixin^-/-^ and LOK^-/-^/SLK^-/-^ cells to about 1.2, indicating greater distortion/waviness of the interface. As with loss of microvilli, loss of just ezrin or LOK is more severe than loss of just radixin or SLK, and both double mutants exhibit the most severe phenotype. This tortuosity of the junctions was rescued by expressing ezrin in the ezrin^-/-^ radixin^-/-^ cells and LOK in the LOK^-/-^ SLK^-/-^ cells (Figure 3A).

**Figure 3:**
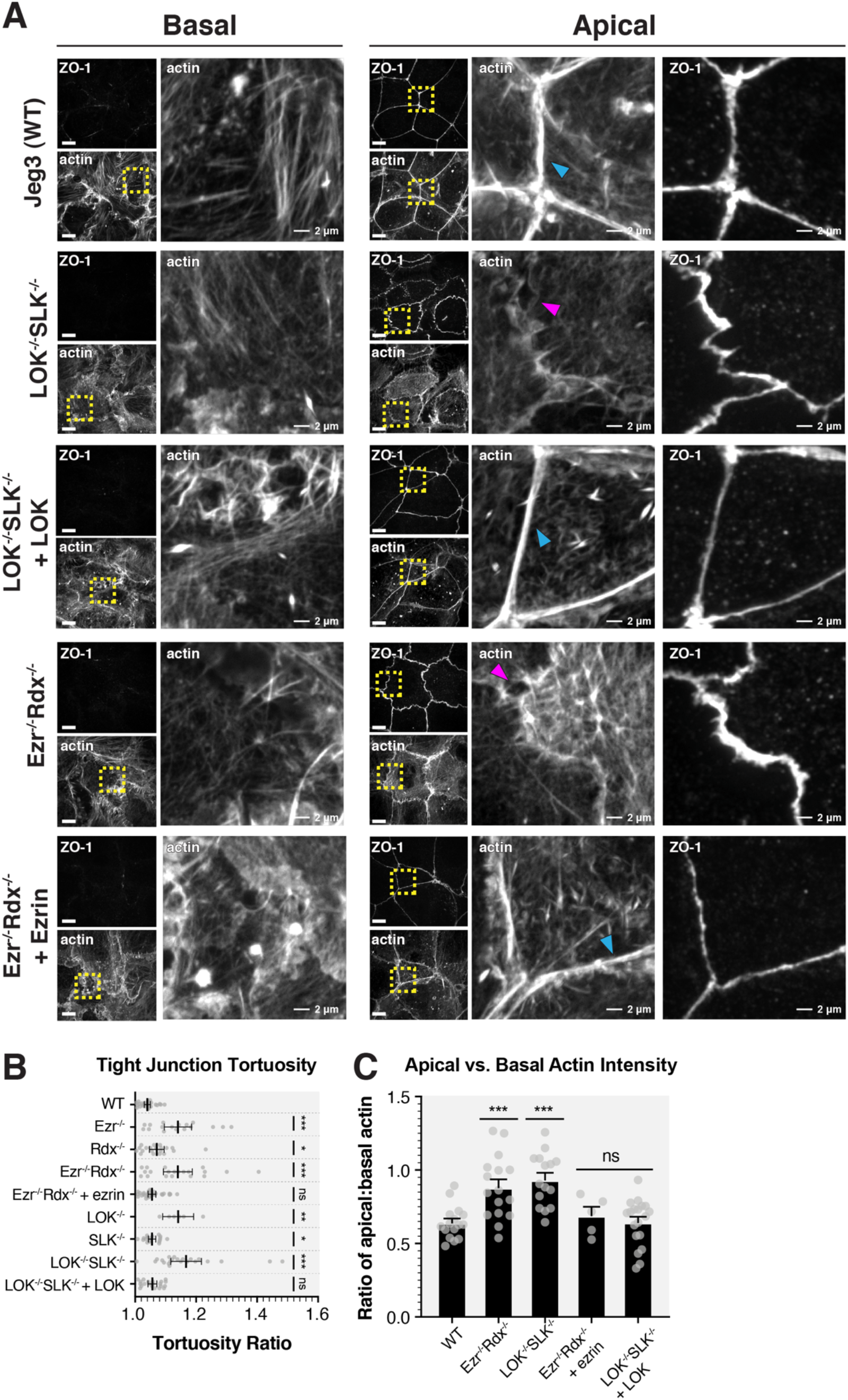
Absence of activated ERM proteins induces the formation of apical actin bundles, alters cell-cell junctions, and compromises the apical rigidity of epithelial cells. (A) Comparison of ZO-1 and actin between basolateral and apical confocal slices. Apical Z-slices were determined by the presence of ZO-1. Blue arrows indicate actin present at cell junctions. Magenta arrows point to actin breaks between neighboring cells. Scale 10μm unless otherwise noted. (B) Quantification of tight junction tortuosity using ZO-1 staining. Each point represents the tortuosity from one tight junction intersection to the next intersection. Center lines respresent mean±SEM. Non-significant (ns) p-values are as follows: Ezr ^-/-^ Rdx ^-/-^ + ezrin = 0.0578 ; LOK ^-/-^ SLK ^-/-^ + LOK = 0.0573. (C) The ratios of relative mean actin intensity values per cell between apical and basolateral cross-sections. Bars represent mean±SEM. Non-significant p-values are as follows: Ezr ^-/-^ Rdx ^-/-^ + ezrin = 0.4188 ; LOK ^-/-^ SLK ^-/-^ + LOK = 0.9555. P-values were calculated with Welch’s t-test (* p≤0.05, ** p≤0.01, *** p≤0.001).

We also noticed an additional striking effect on the actin distribution in which significantly more F-actin is seen spanning the apical domain in the double knockout cells compared to their wild type counterpart. Therefore, we examined the F-actin distribution in the basal and apical regions of the cells. ZO-1 localization is found near the apical region of both wildtype and knockout cells, so we split confocal planes into those containing ZO-1 staining and above (the apical domain), and those below ZO-1 staining (the basolateral domain). Whereas the wildtype cells display actin in microvilli and along the cell junctions, both ezrin^-/-^ radixin^-/-^ and LOK^-/-^/SLK^-/-^ cells were noticeably different with actin bundles extending across the apical domain that correlate with the uneven contour of the tight junctions (Figure 3A). In contrast, there is no apparent difference in the basal actin network between the cells (Figure 3A), which is underscored by the similarity of focal contacts at the basal surfaces (Figure S2A). We quantified this difference by measuring the ratio of the actin intensity at the apical versus the basal side of the cells (Figure 3C). Both KO cells were found to have an increase in the density of actin found exclusively at the apical surface compared to wildtype cells (Figure 3A and C). The aberrant apical actin bundles are rescued by introduction of ezrin into ezrin^-/-^ radixin^-/-^ and of LOK into LOK^-/-^/SLK^-/-^ cells (Figure 3 A-C).

The apical actin structures seen in ezrin^-/-^ radixin^-/-^ and LOK^-/-^ SLK^-/-^ cells contained bundles, similar in appearance to stress fibers. The expression level of the stress fiber marker α-actinin and focal adhesion marker vinculin were unchanged in the knockout cells (Figure S3). We next explored if these bundles could have contractile properties by staining for non-muscle myosin-IIB (myo-IIB). While striated bundles of myo-IIB and actin were not completely absent in the apical region of wildtype cells, a stronger intensity and more frequent clusters of myo-IIB was observed in ezrin^-/-^ radixin^-/-^ cells (Figure 4A, arrows). This phenotype was also seen and enhanced in LOK^-/-^SLK^-/-^ cells where the entire apical domain appeared covered in myo-IIB puncta and actin filaments. Using deconvolution microscopy, we could see characteristic myosin-II sarcomeric-like striations indicative of a contractile structure (Figure 4B). Together our results show that lack of activated ERMs, either due to loss of the proteins or their activating kinases, selectively redistributes actin in the apical domain into contractile bundles.

**Figure 4:**
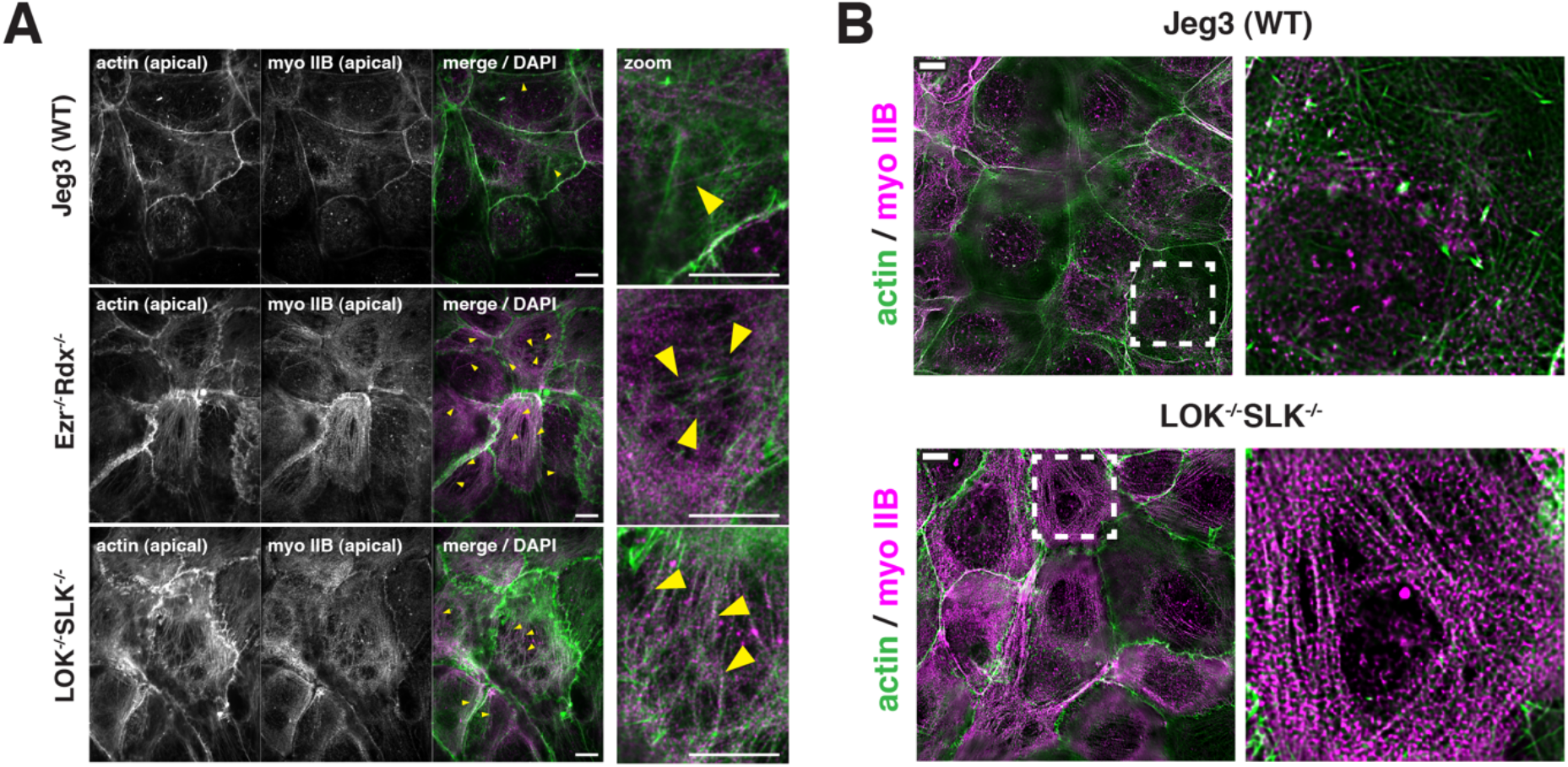
A dense actomyosin network forms at the apical surface of ERM and LOK/SLK knockout cells (A) Maximum projection of immunofluorescence images of the apical confocal Z-slices (2.8μm or 10 confocal slices) indicating actin and non-muscle myosin IIB localization. Areas with contractile fibers are indicated by yellow arrows. (B) Similar apical max projection from deconvolution imaging of the same samples as in (A) showing numerous non-muscle myosin puncta along the apical actin bundles in LOK/SLK knockout cells. Scale bars 10μm.

### Enhanced actin assembly in the apical domain is responsible for the junctional defects of knockout cells

In wildtype cells, apical actin in the microvilli treadmills continuously resulting in turnover in the ~2-10 min time-frame (Loomis et al., 2003; Meenderink et al., 2019). We investigated the possibility that the abnormal actin bundles seen in knockout cells might arise in part from reduced actin turnover in the apical domain. To test this, we treated wildtype cells with the actin stabilizing drug Jasplakinolide to reduce actin turnover. Remarkably, treatment of wildtype cells with 500nM Jasplakinolide for 30 minutes induced both the formation of actin bundles in the apical domain and increased the tortuosity of the tight junctions (Figure 5A and B, lanes 1 and 2) in a manner closely resembling the phenotype of ezrin^-/-^ radixin^-/-^ or LOK^-/-^ SLK^-/-^ cells (Figure 3A). The addition of Jasplakinolide in all three cell lines further increased the abundance of apical F-actin networks across each cell, correlating with the increased tight junction tortuosity and breaks between neighboring cells (Figure 5A, magenta arrows and C).

**Figure 5:**
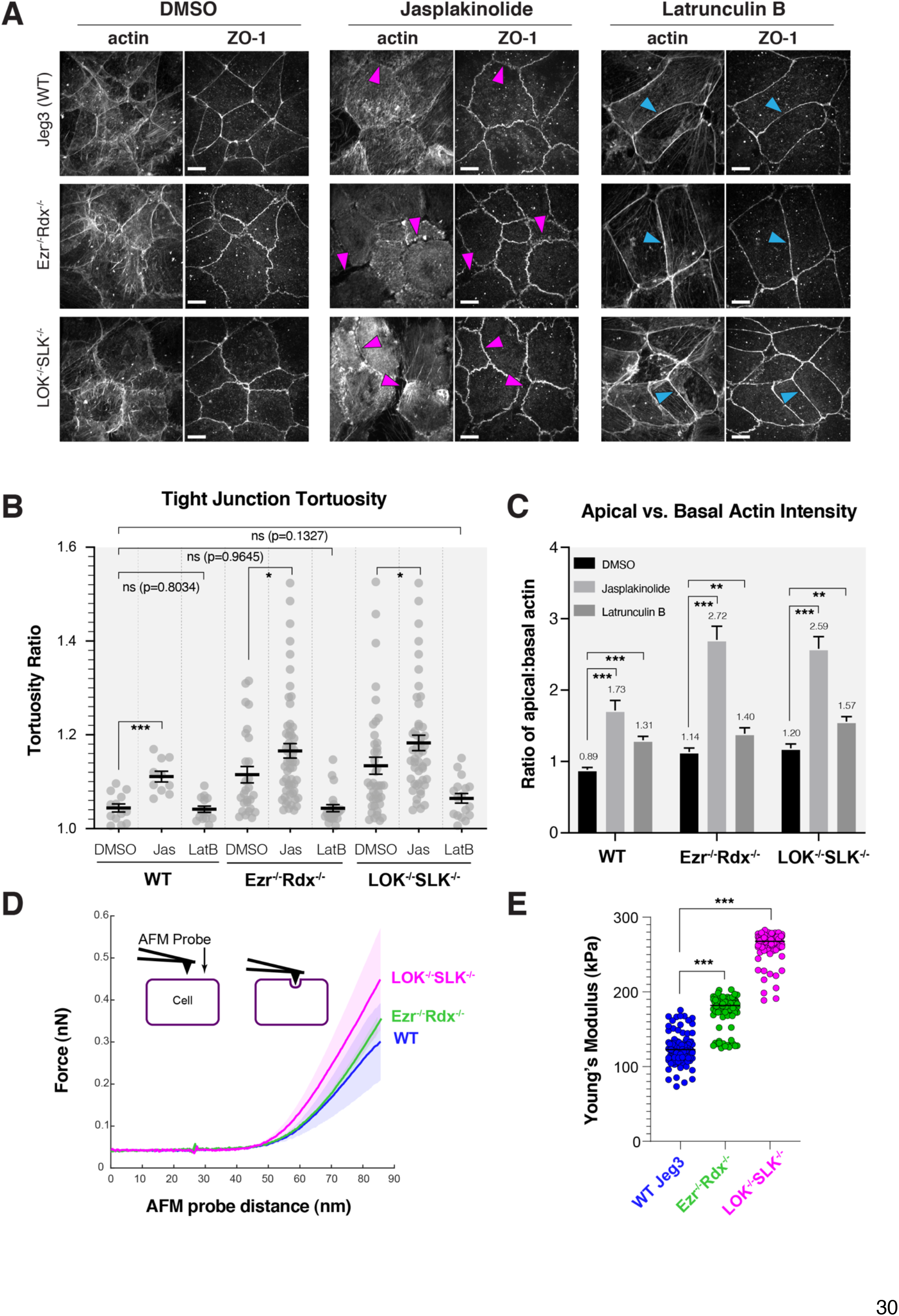
Increased actin polymerization in ERM or LOK/SLK knockout cells leads to junctional defects and mechanical stiffening of the cell. (A) Jeg3 cells treated with DMSO, 500nM Jaspakinolide or 100ng/mL Latrunculin B for 30 min before immunostaining with ZO-1 and actin. Magenta arrows point to actin gaps at junctions while blue arrows represent actin present at cell junctions. Scale bars, 10 μm. (B) Quantification of tortuosity between tight junction markers as described in Figure 3. Jasplakinolide treatment increases tortuosity values, while Latrunculin B rescues tortuosity to wildtype levels. N ≥ 10 cells per condition. Lines represent mean±SEM. P-values were calculated with Welch’s t-test (* p≤0.05, ** p≤0.01, *** p≤0.001). (C) Comparison of actin levels between apical and basolateral regions after treatment with drugs as visualized in panel A. N ≥ 21 cells per condition. Bars show mean±SEM. (D) Averaged force indentation curves for WT Jeg3 (blue), ezrin^-/-^radixin^-/-^ (green) and LOK^-/-^ SLK^-/-^ (magenta); semi-transparent area around each line represents the SEM of the data. A steeper curve indicates a stiffer cell. (E) Young’s modulus stiffness parameters; black line indicates each condition’s mean value (WT N=356, ezrin^-/-^radixin^-/-^ N=304, LOK^-/-^ SLK^-/-^ N=431). Both KO conditions were significantly stiffer than WT cells (*** p=<0.0001, K/S Test).

If enhanced actin assembly in the double knockout cells is responsible for the observed tortuosity of the junctions, increasing depolymerization in these cells might rescue this phenotype. Upon treatment of either of the double-knockout cells with the actin-depolymerizing drug Latrunculin B or actin plus-end-capping drug Cytochalasin D, tight junction tortuosity ratios in the knockout cells were restored to levels comparable to wildtype cells (Figure 5A, B and S5A). Interestingly, Latrunculin B treatment of cells did not completely restore the balance of F-actin between the apical and basolateral domains to wild type levels (Figure 5C). Analysis of the apical/basolateral actin distribution after these drug treatments was complicated by the finding that apical actin appears to be more resistant to disassembly by Latrunculin B than its basolateral counterpart (Figure S5B). Nonetheless, in contrast to Jasplakinolide-treated cells, Latrunculin B treatment results in reorganization of actin towards junctions Figure 5A, blue arrows) and relief of junctional defects (Figure 5B). Meanwhile, stabilization by Jasplakinolide redistributes actin away from the junctions and across the cell promoting junctional defects (Figure 5A and B). In summary, our data shows ezrin^-/-^ radixin^-/-^ and LOK^-/-^ SLK^-/-^ cells become more similar to wildtype when treated with Latrunculin B, while wildtype cells become more like the knockout cells when treated with Jasplakinolide. Therefore in the absence of active ERMs, excessive actin assembly generates contractile bundles that provide forces perpendicular to junctions.

To assess how the actin redistribution might affect the mechanics of the apical surface, we utilized Atomic Force Microscopy (AFM) to measure their stiffness. Cells were allowed to grow to a confluent monolayer and then indented to a maximum force of 1nN (Figure 5D inset). The resulting force vs. distance curves were then used to identify the initial AFM probe tip contact point, allowing us to fit to a hertz equation to calculate the Young’s modulus (E) stiffness parameter (Huth, Sindt, & Selhuber-Unkel, 2019). Indentations from each condition were then averaged to produce a mean curve for wild type, ezrin^-/-^ radixin^-/-^ and LOK^-/-^ SLK^-/-^ cells (Figure 5D). Of the 1,091 total force indentations preformed, both LOK^-/-^ SLK^-/-^ (E = 260±45 kPa Median±) and ezrin^-/-^ radixin^-/-^ (E = 166±52 kPa) cells are significantly stiffer then WT cells (E = 117±65 kPa) (Figure 5E). The finding that LOK^-/-^SLK^-/-^ cells are more rigid than ezrin^-/-^ radixin^-/-^ cells is consistent with the enhanced levels of stress-fiber-like bundles seen in the apical domain of LOK^-/-^SLK^-/-^ cells described in Figure 4. Together these results indicate that activated ERM proteins suppress the formation of apical actin bundles to affect the mechanical properties of the apical domain.

### Activated ezrin Rescues both ezrin^-/-^ radixin^-/-^ and LOK^-/-^SLK^-/-^ cells

The phenotypic similarity of the ezrin^-/-^ radixin^-/-^ and LOK^-/-^ SLK^-/-^ cells, suggested that essentially all the phenotypes of LOK^-/-^ SLK^-/-^ cells are due to the lack of ERM phosphorylation. If this is the case, introduction of mutationally activated ezrin in either of the double knockout cells should rescue them in a similar manner. We therefore introduced the constitutively active phosphomimetic ezrin-T567D mutant into both ezrin^-/-^ radixin^-/-^ and LOK^-/-^/SLK^-/-^ cells. Remarkably, phosphomimetic ezrin suppressed both equally with restoration of microvilli (Figure 6A). The rescue isn’t perfect, as constitutively active ezrin is seen both apically and basolaterally because restriction to the apical domain requires ezrin phosphocycling (Viswanatha et al., 2012). Expression of constitutively active phosphomimetic ezrin-T567D is also able to suppress tight junction defects and the excess apical actin bundles seen in knockout cells implying that active ezrin can regulate actin at the cortex (Figure 6B-D). These results suggest that the primary role for LOK and SLK in Jeg3 cells is to phosphorylate ERMs.

**Figure 6:**
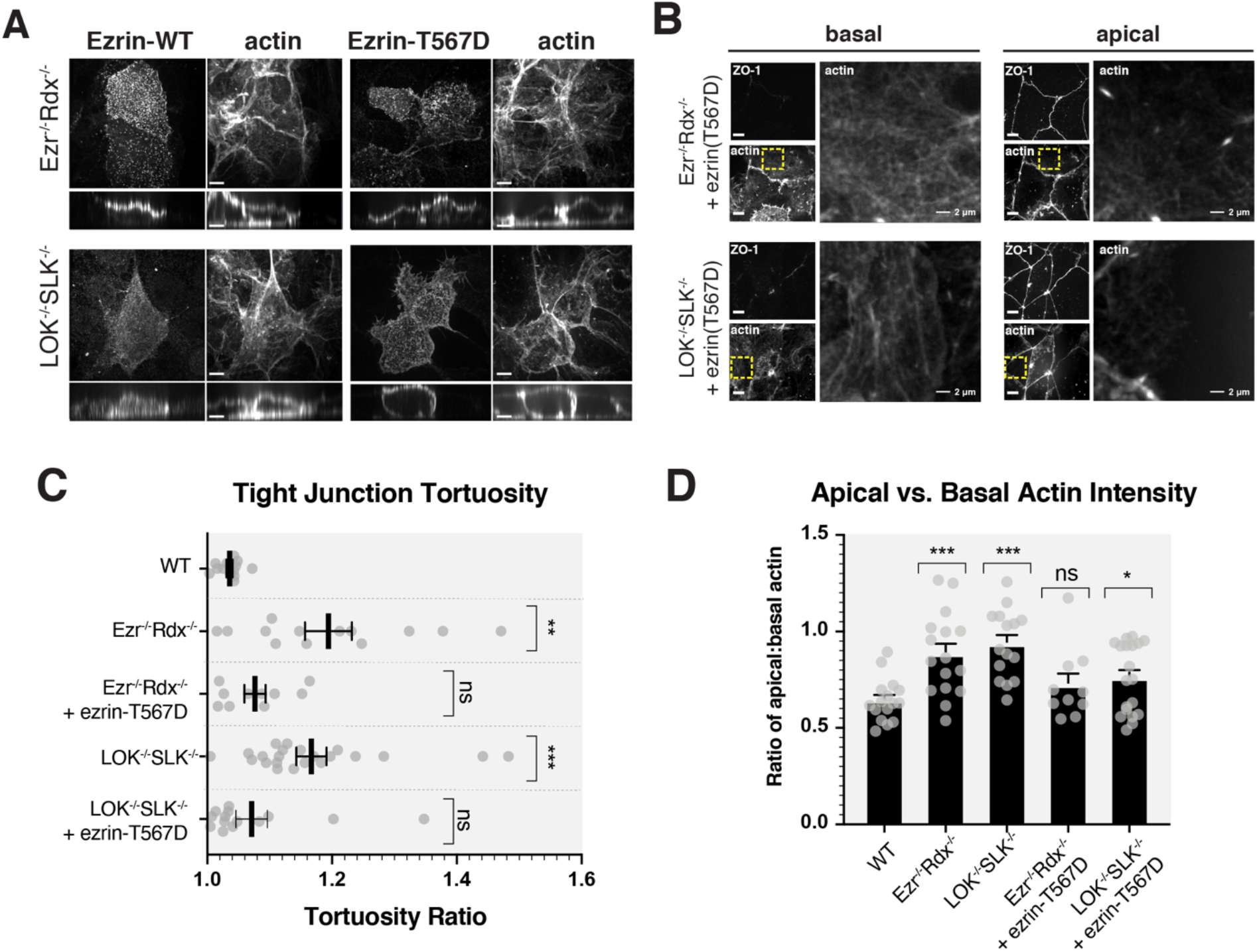
Constitutively active ezrin rescues apical actin distribution in knockout cells. (A) Jeg3 knockout cells expressing wildtype ezrin or T567 phosphomimetic were stained for ezrin and actin. Maximum intensity projections and vertical cross-sections are shown. (B) Comparison of ZO-1 and actin between basolateral and apical confocal slices between Jeg-3 knockout cells expressing phosphomimetic ezrin-T567D. Apical Z-slices were determined by the presence of ZO-1. Yellow boxes indicate the region of the image that was expanded on the right. (C) Quantification of tight junction tortuosity using ZO-1 staining. N ≥ 10 cells per condition. Center lines show mean±SEM. Non-significant (ns) p-values are as follows: Ezr ^-/-^ Rdx ^-/-^ + ezrin-T567D = 0.0623; LOK ^-/-^ SLK ^-/-^ + ezrin-T567D = 0.2445 (D) The ratios of relative mean actin intensity values per cell between apical and basolateral cross-sections. N ≥ 21 cells per condition. Nonsignificant p-value for Ezr ^-/-^ Rdx ^-/-^ + ezrin-T567D = 0.4188. Bars show mean±SEM. P-values were calculated against wildtype with Welch’s t-test (* p≤0.05, ** p≤0.01, *** p≤0.001). Scale bars, 10μm; unless otherwise noted.

### Phosphorylated ERM proteins regulate myosin-II activity through RhoA Activation

The appearance of apical stress-fiber-like cables in cells lacking activated ERMs is suggestive of local enhanced RhoA activity. Rho-associated kinase (ROCK) is a major effector of RhoA-GTP that can activate myosin-II by directly phosphorylating myosin light chain 2 (MLC2), and this is counteracted by the phosphatase PP1 utilizing the Mypt1 subunit. Total non-muscle myosin-II expression showed no differences between wild type and LOK^-/-^SLK^-/-^ or ezrin^-/-^ radixin^-/-^ cells (Figure S2B). The level of endogenous MLC2 phosphorylation was found to be very low in wildtype cells and 3-4 fold enhanced in both ezrin^-/-^ radixin^-/-^ and LOK^-/-^SLK^-/-^ cells (Figure 7A and B). To increase the level of phosphorylation, we treated cells briefly with calyculin A that inhibits the phosphatase PP1. Treatment of wildtype cells with calyculin A enhanced the level of MLC2 phosphorylation, and greatly enhanced the level seen in both of the knockout lines. These results reveal that activated ERM proteins negatively regulate phosphorylation of MLC2 and hence myosin-II activation.

**Figure 7:**
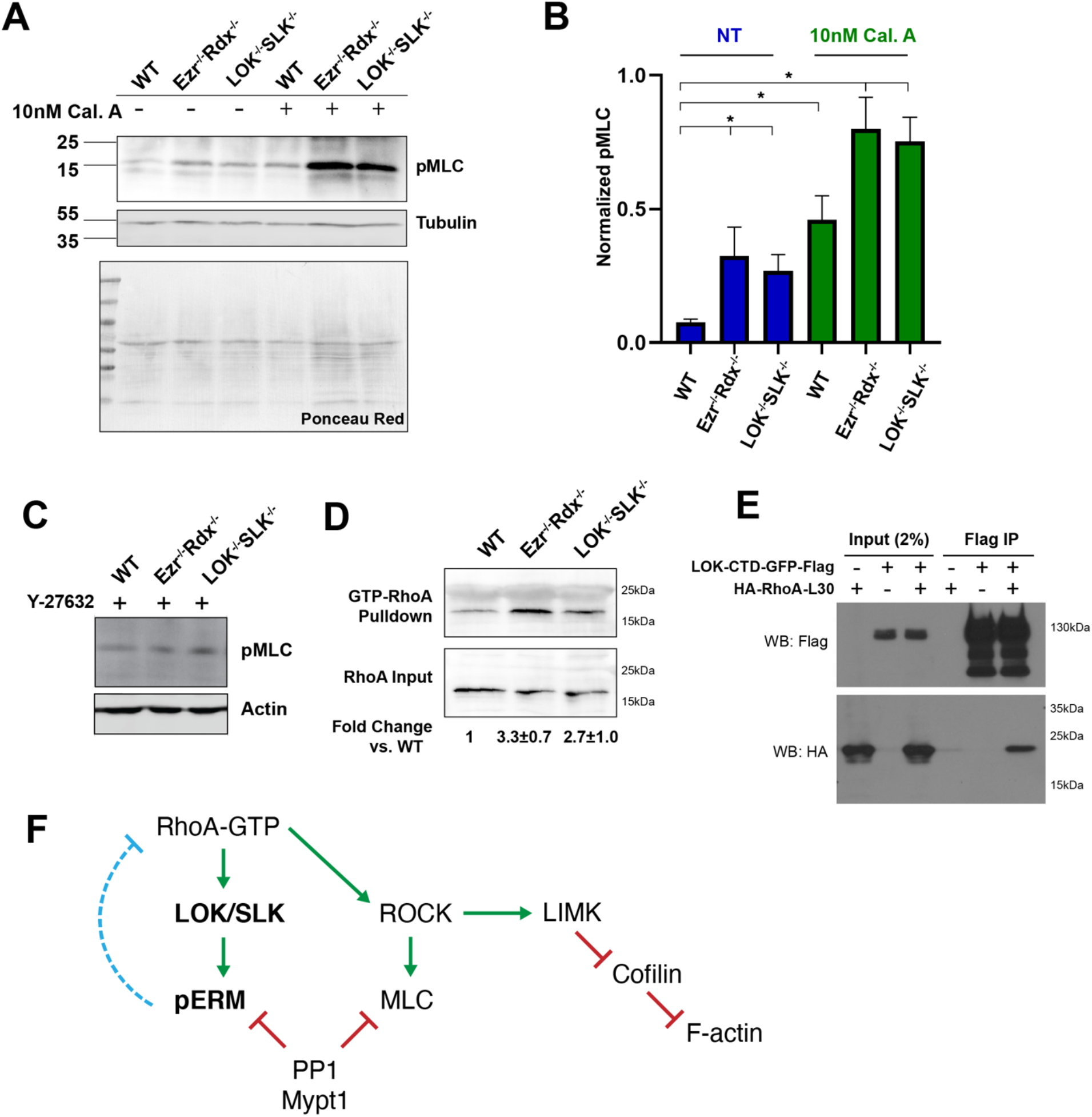
Phosphorylated ERMs negatively regulate myosin activation through RhoA. (A) Phospho-myosin light chain (pMLC) (T18/S19) western blot with tubulin and total protein staining to indicate equal loading. (B) Quantification of pMLC staining normalized to loading control (bars show mean±SEM; control:N = 4; 10 min of 10nM CalA: N = 3). (C) Western blot of pMLC with actin loading control in Jeg3 cells first treated with ROCK inhibitor Y-27632 showing reversal of the pMLC hyperphosphorylation phenotype in KO cells. (D) Representative Western blot of active RhoA-GTP pull down results. Upper row: Rho-A blot of the Rhotekin beads following pulldown, Lower row: blot of Rho-A of 4% of input lysate. Normalized fold change of the RhoA pull down band intensity compared to WT (mean±SEM; N = 5). (E) LOK-CTD-GFP-Flag pulldown of cells expressing constitutively active RhoA (RhoA-L30) indicating that C-term LOK binds active RhoA. (F) Model of Rho A signaling negatively regulated by phospho-ERMs (dotted blue line).

MLC2 phosphorylation is either mediated by ROCK, or via Ca^2+^-calmodulin-dependent myosin light chain kinase. To explore which pathway is involved, we treated wildtype and double knockout cells with the specific ROCK inhibitor Y-27632. This resulted in equalization of phospho-MLC2 levels (Figure 7C), indicating that the enhanced level of phospho-MLC2 seen in knockout cells is upstream of ROCK. We therefore measured the level of endogenous RhoA-GTP by passing total cell lysates over Rhotekin-GST beads and measuring the relative levels of RhoA retained. Both ezrin^-/-^ radixin^-/-^ and LOK^-/-^SLK^-/-^ cells were found to have about a 3-fold higher level of active RhoA than wildtype cells (Figure 7D). In addition to regulating MLC2 phosphorylation, ROCK also regulates actin dynamics through the LIM kinase-cofilin pathway (Arber et al., 1998; Maekawa et al., 2017; Yang et al., 1998), which may explain in part the altered actin dynamics discussed earlier.

A recent report showed that active RhoA binds directly to the C-terminal domain of SLK and this promotes its dimerization and ability to phosphorylate ERM proteins (Bagci et al., 2020). In earlier work we showed that expression of the C-terminal region of LOK (LOK_310-968-GFP-Flag or LOK-CTD-GFP-Flag) acts as a potent dominant negative to strongly inhibit phosphorylation of ERM proteins (Viswanatha et al., 2012). To explore if this region of LOK binds to RhoA, we expressed LOK-CTD-GFP-Flag either alone or with constitutively active HA-RhoA-L30 and immunoprecipitated with Flag antibodies in WT Jeg3 cells (Figure 7E). RhoA was recovered in the Flag immunoprecipitates, indicating an interaction between the two. Therefore, both LOK and SLK are effectors of RhoA.

Collectively these results suggest a model in which RhoA selectively regulates the apical domain of epithelial cells in a negative feedback loop involving active ERM proteins (Figure 7F). RhoA activates both the kinases LOK/SLK and ROCK to mediate phosphorylation of ERM proteins and MLC2, respectively. ROCK also negatively regulates PP1, the phosphatase that dephosphorylates both pMLC and pERMs. Phosphorylated ERMs localized exclusively in the apical domain regulate RhoA in a negative feedback loop.

### Phosphorylated ERMs negatively regulate myosin contractility

If the model presented in Figure 7F is valid, a prediction is that a functional difference in contractile force production should exist between wildtype and knockout cells. It is known that cultured cells treated with calyculin A to elevate the level of myosin-II activity will ultimately contract (Ishihara et al., 1989). Therefore, we examined the contraction of spread LOK^-/-^SLK^-/-^ and ezrin^-/-^radixin^-/-^ cells compared with wildtype cells in the presence of 10nM Calyculin A over the course of 1 hour. Under this treatment a small fraction of WT cells showed rounding and detachment from the plate. In a dramatic difference, LOK^-/-^SLK^-/-^ and ezrin^-/-^radixin^-/-^ cells showed significant rounding and detachment from neighboring cells in under 30 minutes (Figure 8A, Movie S1). We confirmed that the contractility is a result of enhanced myosin-II activity as the calyculin A induced contraction was prevented by inclusion of the myosin-II inhibitor blebbistatin, which showed no rounding or contraction in any cell type after 1 hour (Figure 8B, Movie S2). These results are in agreement with Figure 7A showing an increased level of MLC phosphorylation in the knockout cells in the presence of calyculin A. However, calyculin A also results in enhanced phosphorylation of ERMs (Viswanatha et al., 2012), so we wished to examine if the contractility difference induced by calyculin A was present in cells where the level of active ezrin was unchanged. To achieve this, we stably expressed the active phosphomimetic ezrin-T567E in double knockout cells. Under these conditions both LOK^-/-^ SLK^-/-^ and ezrin^-/-^ radixin^-/-^ cells expressing ezrin-T567E the contraction induced by calyculin A was greatly reduced (Figure 8B, Movie S3). This implies that it is the loss of active ERMs, and not the loss of LOK and SLK independent of ERMs, that regulates the contractility due to enhanced RhoA-GTP.

**Figure 8:**
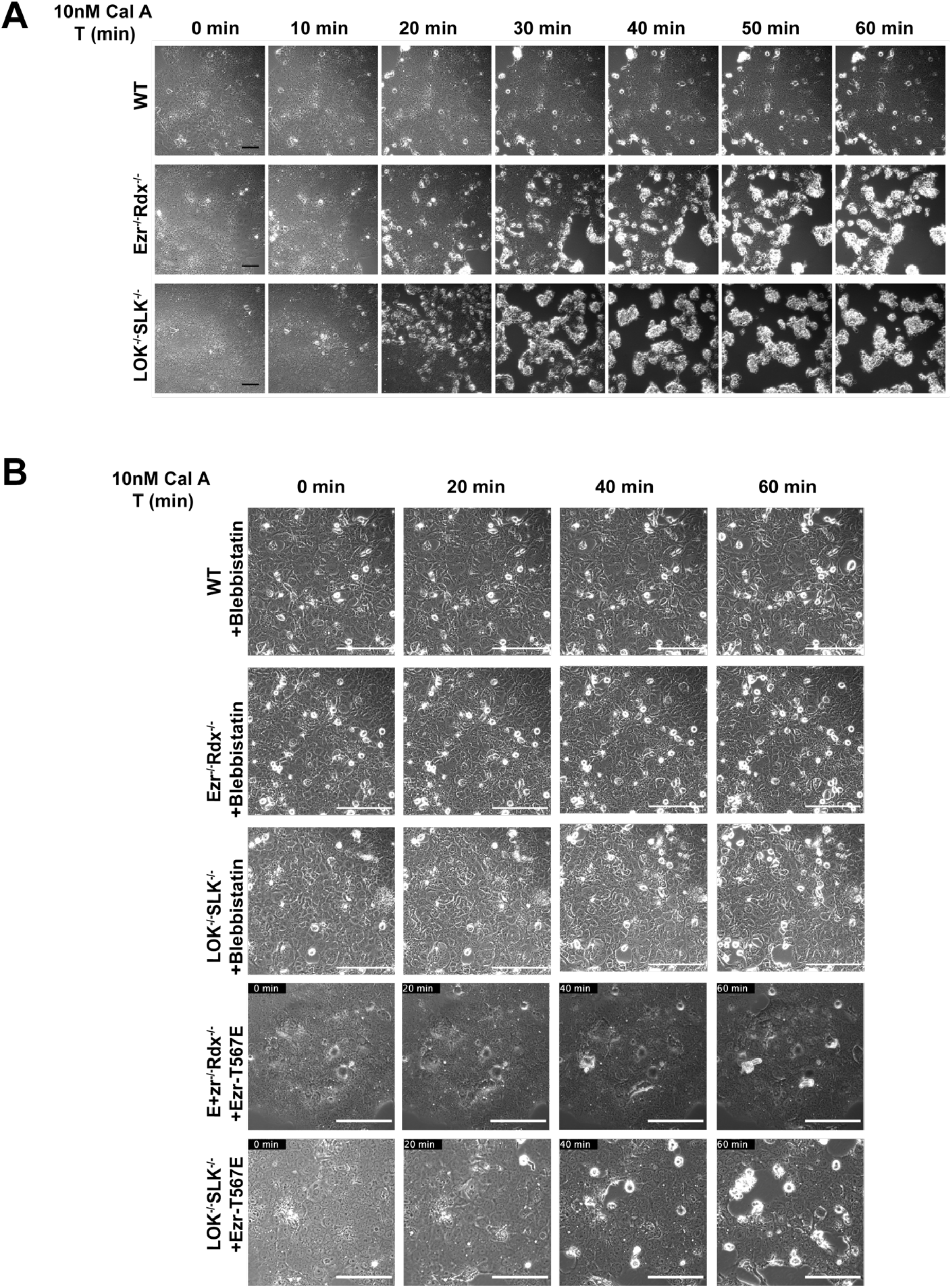
Phosphorylated ERMs negatively regulate myosin contractility. (A) Representative phase contrast still images from movies (S1) showing the effect of 10nM calyculin A on WT or KO Cells. (B) Same as (A) except for 30 min treatment with 25uM blebbistatin (Movie S2), or transfection to express phosphomimetic ezrin-T567E (Movie S3), prior to calyculin A treatment. Scales 100μm.

## Discussion

In this study we have investigated the phenotypes conferred by loss of ERM proteins, or their activators, LOK and SLK. In some cells loss of LOK and SLK, or in fact just ezrin in HeLa or Caco-2 cells, are too unhealthy to maintain, indicating that ERM proteins and their activating kinases can be almost essential. This is consistent with the situation in mice, flies and the nematode worm, where loss of ezrin or the single ERM protein (flies and worm), is lethal (Jankovics et al., 2002; Saotome et al., 2004). Additionally, the loss of SLK in mouse or the single homolog Slik in the fly is also lethal (Hipfner and Cohen, 2003). Loss of just LOK in the mouse has a more modest phenotype (Belkina et al., 2009), presumably due to partial compensation by SLK. We were fortunate to start with a cell line, Jeg-3, that can tolerate loss of either all ERM proteins, or both LOK and SLK, as this allowed us to study their phenotypes. As far as we are aware, these are the first vertebrate cells isolated genetically lacking either all ERM proteins or the kinases LOK and SLK.

Previous studies have shown that wildtype epithelial cells maintain ~50% of their ezrin in the active, phosphorylated state with cycling between active and inactive states occurring on the scale of ~2 minutes (Viswanatha et al., 2012). Proper preservation of this balanced system is critical for maintaining a polarized morphology (Viswanatha et al., 2012) and is consistent with mislocalization, overexpression, and hyper-phosphorylation of ezrin found in types of human cancers (reviewed in Clucas and Valderrama, 2015). Our finding that loss of LOK and SLK results in ablation of all detectable ezrin-T567 phosphorylation is consistent with LOK and SLK being the major, if not only, kinases that can phosphoryate the ERM regulatory threonine (T567 in ezrin). This result is in agreement with earlier descriptions that Slik is solely responsible for the activation of fly moesin (Hipfner et al., 2004).

A remarkable aspect of our results is that cells lacking either all ERM proteins or LOK and SLK are phenotypically very similar. Both grow slower, have lost all apical microvilli, have greatly reduced junctional actin but also an aberrantly high level of apical F-actin and myosin, have wavy cell junctions, have a stiffer apical domain, contract abnormally in the presence of the phosphatase calyculin A, and both have elevated levels of Rho-GTP. This raises the question whether the ERM proteins are the sole substrate of LOK/SLK. An intricate multi-step mechanism is involved in phosphorylation of ezrin by LOK. In outline, it requires priming of ezrin by PIP2, then insertion of the C-terminal region of LOK between the ezrin FERM and C-terminal F-actin binding domain, which gives access for the kinase domain to bind a recognition site and ultimately phosphorylate T567 (Pelaseyed et al., 2017). This mechanism, together with the strong preference for a tyrosine two residues upstream of the targeted threonine (Belkina et al., 2009), makes phosphorylation of ERM proteins highly selective. This elaborate coincidence detection mechanism involved in LOK phosphorylation of ezrin coupled with the ability of mutationally active ezrin (ezrin-T567D) to suppress the phenotype of LOK/SLK cells strongly supports the notion that ERM proteins are the major functional substrates. Using a phospho-proteomics approach in unpublished work, we sought to identify additional LOK/SLK substrates. While we encountered many phosphopeptides whose level was elevated in wild type cells compared with LOK/SLK knockout cells, none had the appropriate LOK/SLK consensus sequence. Therefore, LOK/SLK and ERM proteins appear to work together as a functional unit. Although we were not able to identify additional LOK/SLK substrates, technical limitations may have obscured them from our analysis. In support of this possibility, the phenotypes of LOK/SLK cells enumerated above were almost always more severe than in the ERM knockout cells. Thus, the possibility remains that there is another minor substrate, functionally redundant with ERM proteins, cannot be ruled out.

While we expected to see loss of microvilli in cells lacking ERM proteins, we were surprised to encounter an extensive actin/myosin-II network in the apical domain replacing the junctional and microvillar actin. This phenotype was also seen in LOK/SLK knockout cells, implying that active ERM proteins can regulate the myosin and actin distribution in the apical domain. RhoA-GTP is known to positively regulate non-muscle myosin-II activity through ROCK1 as well as F-actin turnover through LIM kinase and cofilin. Since both actin turnover and myosin contractility are altered in the absence of pERMs, we then considered a possible over-activation of RhoA in the knockout cells. Indeed, the level of RhoA-GTP was significantly elevated in both ERM and LOK/SLK knockout cells. Thus, active ERMs are negative regulators of RhoA (Figure 7F).

There have been several indications of a connection between ERM protein function and RhoA. First, ROCK1 was reported as the kinase that phosphorylates ERM proteins, in part because ROCK1 over expression resulted in formation of microvilli in cos-7 cells and its reported ability to phosphorylate radixin *in vitro* (Matsui et al., 1998; Oshiro et al., 1998). With the finding that Mypt1/PP1 is the phosphatase responsible for dephosphorylating ERM proteins (Fukata et al., 1998), and that Mypt1/PP1 is negatively regulated by ROCK1 (Velasco et al., 2002), the earlier result can be explained. In a genetic study in the fly, phenotypes resulting from loss of moesin can be rescued by also reducing the level of RhoA (Speck et al., 2003). Recent work has shown that RhoA-GTP directly dimerizes and activates SLK (Bagci et al., 2020). As phosphorylated ERM proteins act as negative regulators of RhoA-GTP, a local feedback cycle exists in which RhoA-GTP and pERMs regulate each other’s activity specifically across the apical domain (Figure 7F).

RhoA is a well recognized spatial regulator of the adherens junctions between epithelial cells (reviewed in Hartsock and Nelson, 2008; Marjoram et al., 2014; McCormack et al., 2013). Active RhoA is highly enriched at junctions through the action of various regulators, including the RhoA-GEFs p114RhoGEF and ECT2, and indirectly through active myosin IIA recuitment of ROCK1 to phosphorylate and inactivate the ability of Rnd3 to recruit p190B RhoA-GAP (Ratheesh et al., 2012; Reyes et al., 2014; Terry et al., 2011). Misregulation of RhoA can affect the mechanical tension between junctions of neighboring epithelial cells (Zihni et al., 2014). Indeed, this is what we observe by knocking out active ERMs. As a consequence of removing active ERMs, we find that F-actin bundles are re-distributed away from the junctions and into actomyosin networks across the apical terminal web. This state can be mimicked in wildtype epithelial cells by introducing an actin stabilizing drug like Jasplakinolide. Conversely, the actin in knockout cells can be re-directed back towards junctions by adding actin depolymerizing drug Latrunculin B. A similar phenomenon is seen when anillin is manipulated in epithelial cells, where anillin knockdown cells produce low tensile forces and overexpression produced high tensile forces due to misregulation of medial-apical F-actin (Arnold et al., 2019). Thus, local spatial regulation of proteins like ERMs and anillin can influence the RhoA-dependent actin turnover at the apical domain and junctions of epithelial cells. As LOK is an effector of active RhoA and is located to the microvilli, and not the cell-cell junctions, a subpopulation of RhoA must be specifically regulated at across the apical surface of epithelial cells. The activity of this local RhoA subpopulation is negatively regulated by phosphorylated ERM proteins, presumably through either a Rho-GEF or a Rho-GAP. In the fly RhoGAP Conundrum has been shown to bind moesin and act as a negative regulator of Rho1. However, since deletion of Conundrum has no obvious phenotype there is likely functional redundancy with another RhoGAP (Neisch et al., 2013). Our future work will be aimed at identifying the factor(s) that allow active ERMs to mediate local negative regulation of RhoA-GTP.

A more distant member of the ERM protein family is merlin, the product of the tumor suppressor gene *(NF2)* in which defects result in Neurofibromatosis-II (Sanson et al., 1993; Trofatter et al., 1993). Merlin has an N-terminal FERM domain and a negative regulatory C-terminal. The FERM domain of merlin can bind the C-terminal domain of ERM proteins, and *vice versa* (Bretscher et al., 2000). Considerable work suggests that ERM proteins and merlin act antagonistically, especially in the organization of the plasma membrane during endocytosis (Chiasson-Mackenzie et al., 2018; Hebert et al., 2012). It will be interesting to explore if active merlin can regulate RhoA activity, or if merlin-deficient cells that fail to modulate the activity of ERM proteins have a lower level of RhoA-GTP.

In summary, we have found that ERM proteins and their activators LOK/SLK function in the same pathway as a unit to build microvilli on the apical surface of epithelial cells and locally modulate the level of RhoA-GTP in a feedback inhibition cycle. The next challenge will be to understand how this feedback loop is regulated and restricted to the apical domain.

## Materials and Methods

### Reagents & cDNAs

Ezrin antibodies were either a mouse anti-ezrin antibody (CPCT-ezrin-1 supernatant concentrate obtained from the Developmental Studies Hybridoma) used at 1:1,250 (Western blot) or 1:100 (immunofluorescence) or a previously characterized rabbit polyclonal antibody raised against full-length human ezrin (Bretscher, 1989) and used at 1:1,000 (Western blot) or 1:200 (immunofluorescence). Rabbit anti-EBP50 was also a previously characterized antibody (Reczek et al., 1997) and was used at 1:50 (immunofluorescence). Mouse anti-ZO1 (BD Biosciences, Cat #610966) was used at 1:100 (immunofluorescence). Rabbit anti-radixin (Cell Signaling, Cat #C4G7) used at 1:1000 (Western blot) to blot for both radixin (80kDa) and moesin (75kDa). Rabbit anti-pERM (raised against re-combinant phosphopeptide) was used at 1:1000 (Western blot). Rabbit anti-LOK used at 1:500 (western blot) and rabbit anti-SLK used at 1:100 (Western blot) were purchased from Bethyl Laboratories, Inc. Mouse anti-Flag used at 1:5,000 (Western blot) and mouse anti-tubulin used at 1:5,000 (Western blot) were obtained from Sigma-Aldrich. Mouse anti-GFP used at 1:100 (immunofluorescence) was obtained from Santa Cruz Biotechnology, Inc. Rabbit antibodies for anti-myosin-light-chain-2 (Cat #3672) and anti-phospho-myosin-light-chain-2 (raised against phospho-Thr18/Ser19; Cat #3674) were purchased from Cell Signaling and used at 1:500 (Western blot). Monoclonal Mouse anti-RhoA antibody (Cytoskeleton inc. Cat#ARH05) was obtained through the Cytoskeleton inc Rho A activation Assay Biochem Kit and used at 1:500. Rabbit antibody for anti non-muscle-myosin-IIB from (BioLegend, Cat. No. 909902) was used at 1:100 (Western blot and immunofluorescence). Rabbit polyclonal antibodies raised by standard procedures against Vinculin, brush boarder myosin-II and alpha-acitinin purified from chicken gizzard were used at 1:2000, 1:200 or 1:200 respectively for supplemental figure 1 western blotting. For actin staining, alexa fluor 488 or 647 phalloidin (Molecular Probes) was used at 1:250. Wheat Germ Agglutinin conjugated to alexa fluor 488 was also purchased from Molecular Probes and used at 1:300 to stain cell membranes.

Phos-tag was purchased from Wako Chemicals. Jasplakinolide (Cat #11705) and Y-27632 (Cat #100005583-5) were purchased from Cayman Chemicals. DMSO (Cat #D2650), Latrunculin B (Cat #L5288) and Cytochalasin D (Sigma Cat #C8273) were purchased from Sigma-Aldrich. Calyculin A (Cat #BML-El92-0100) was purchased from Enzo Life Sciences and Blebbistatin (Cat #B592500) was purchased from Toronto Research Chemicals.

Ezrin point mutants T567E, and T567A were previously generated as described in Viswanatha et al., 2012. Sequences for LOK-GFP-Flag, LOK-CTD-GFP-Flag (or LOK_310-968-GFP-Flag) and LOK-K65R-GFP-Flag were also previously generated in the lab (Pelaseyed et al., 2017; Viswanatha et al., 2012). To generate stable cell lines, ezrin and LOK cDNAs were subcloned into pCDH lentivector (System Biosciences). Puromycin gene in pCDH was then substituted for blasticidin using Gibson assembly. The lentivectors were then transfected with psPAX2 and pCMV-VSVG before virus collection and transduction into Jeg-3 cells. Cells were then grown under blasticidin selection at 5.0 μg/ml for 1-2 weeks prior to immunofluorescent experiments. Stable expression for ezrin was validated using either an ezrin antibody for ezrin constructs or GFP expression for LOK constructs.

### Cell culture

Jeg-3, HeLa and Caco-2 cells (ATCC) were maintained in a 5% CO2 humidified chamber at 37°C. Jeg-3 cells were maintained in MEM with 10% FBS, penicillin/streptomycin and GlutaMax (Thermo Fisher). Cells were cultured on Corning 100mm x 20mm cell culture polystyrene dishes. HeLa and Caco-2 cells were maintained in DMEM with 10% FBS and penicillin/streptomycin. Knockout cell lines were maintained with additional 2.0 μg/ml Puromycin (Sigma) selection. Transient transfections were done using either Lipofectamine 3000 (Invitrogen) according to the manufacturer’s instructions or polyetheylenimine reagent (PolyPlus) as previously described (Viswanatha et al., 2012).

Single guide RNAs (sgRNAs) were designed using CRISPR analysis tools on Benchling and cloned into puromycin-resistant pLenti-CRISPRV2 (AddGene #49535) as described in Sanjana et al., 2014 and Shalem et al., 2014 (Sanjana et al., 2014; Shalem et al., 2014). The following sgRNA sequences were used: 5’ - GCAATGTCCGAGTTACCACCA - 3’ (ezrin), 5’ - AGAAGCAGAACGACTTGAAA - 3’ (radixin), 5’ - GTAAGACTCACCCAGCATGA - 3’ (LOK), 5’ - GCAGTACGAACACGTGAAGA - 3’ (SLK). Each lentiviral construct was then transfected into 293TN cells with psPAX3 and pCMV-VSVG (a gift from Jan Lammerding, WICMB/Cornell, Ithaca, NY) for 48-72 hours before virus collection. Target cells (Jeg3, HeLa or Caco-2) were then transduced with either one or two lentiviruses in order to generate a mixed population of single and double knockout cells. Cells were sorted into single cells and then expanded in puromycin selection before screening by immunofluorescence and Western blotting.

Growth curves of Jeg3 cells were preformed by plating cells on low evaporation lid, flat bottom 96 well plates (Corning Cat. #3595). Once plated, cells were seeded at 3,000 cells per well. Plates were then imaged once per hour for 100hrs using a 20X objective using an Incucyte Zoom v2016 (Essen BioSciences) kept in standard cell incubation chamber conditions. Raw data Images were collected and analyzed using Incucyte. Graphs were assembled and exported using Graphpad/Prism (version 8).

### Western blotting and immunoprecipitation

Western Blot analysis of cell lysates were done using 6%-12% split SDS-PAGE gels while 6% gel was used for phos-tag experiments. Phos-tag reagent was added at a final concentration of 50uM to a standard Tris-glycine-SDS polyacrylamide gel according to the manufacturer’s recommendations. Gels were transferred to a PVDF membrane and blocked with 5% milk in TBS + 0.5% Tween-20. Primary antibodies were incubated with the membrane in 5% bovine serum albumin either for 1 hour at room temperature or overnight at 4°C. Bands were detected with HRP (Thermo Fisher) or infrared fluorescent secondary antibodies (Invitrogen or LI-COR Biosciences). Membranes were imaged using a scanner (Odyssey; LI-COR Biosciences).

For detecting myosin-light-chain and phospho-myosin-light-chain, 16% or split 7.5-17.5% SDS-PAGE gels were used and transferred to PVDF membranes with 0.2um pore size was used (Millipour Immuoblion-P^SQ^). The membrane was then blocked with Immobilon Block-PO phosphoprotein-blocking buffer from EMD-Millipore and incubated with primary antibody solution overnight before developing with chemiluminescent reagents (Radiance Q Plus, Azure Biosystems) on a Bio-Rad ChemiDoc. Relative band intensities were calculated in ImageJ and normalized to a loading control and exported to Graphpad/Prism for statistical analysis. Caylculin A treatments for MLC Blotting were performed by incubating the drug with cells for 10 minutes at 37°C prior to lysis. Lysis of cells for MLC blotting was performed with warm (70°C) Laemmli sample buffer, followed by immediate scraping and boiling.

To determine an interaction between LOK and constitutively active RhoA-L30, cells were transiently co-transfected with LOK-CTD-GFP-Flag and HA-RhoA-L30 (a kind gift from the Cerione lab, Cornell, Ithaca, NY). After 24 hours, cells were lysed and then solubilized in cold immunoprecipitation buffer (25 mM Tris, pH 7.4, 5% glycerol, 150 mM NaCl, 50 mM NaF, 0.1 mM Na3VO4, 10 mM β-glycerol phosphate, 8.7 mg/ml paranitrophenylphosphate, 0.5% Triton X-100, 0.1 μM calyculin A, and protease inhibitor tablet [Roche]) and immunoprecipitated for 2 h using Flag M2 affinity gel (Sigma-Aldrich). Immunoprecipitates were then extensively washed in immunoprecipitation wash buffer (25 mM Tris, pH 7.4, 5% glycerol, 150 mM NaCl, 50 mM NaF, and 0.2% Triton X-100) and then eluted in 200 μg/ml 3×Flag peptide, denatured in Laemmli buffer, resolved by SDS-PAGE, transferred to polyvinylidene difluoride, and developed with HRP western detection.

### Active RhoA pull-down

For GTP-RhoA pull down assay the Rho Activation Assay biochem kit (Cytoskeleton inc, Cat# BK036) was used as described in the product manual. In summary, WT Jeg-3 or KO cell lines were plated and grown for 3 days on Corning 100mm x 20mm cell culture polystyrene dishes. An optimum confluency of 70-80% was used as higher confluences can result in partial loss of a monolayer and disruption of apical morphology in Jeg-3 cells. Cells were lysed using the kit lysis buffer and protease inhibitor cocktail then clarified at 10000 x g at 4°C for 1 min. The lysate was then snap frozen in liquid nitrogen as quickly as possible to reduce RhoA-GTP hydrolysis. Frozen lysates were then stored at −80°C. Protein concentrations were measured using the Bradford reagent and absorption at 595nm. Upon thawing aliquots, lysate protein concentration was then normalized to a uniform protein concentration using kit lysis buffer. Equal concentrations of lysate were then passed on 100ug of Rhotekin-RBD beads. The beads were then washed, pelleted and finally boiled with 20ul of Laemmli sample buffer. Positive and negative controls using WT-Jeg 3 lysate incubated with either GTPys or GDP for 15 minutes prior to passing over the Rhotekin-RBD bead were used to confirm the detectable range of western blot detection, to confirm Rho-A hydrolysis activity and Rhotekin bead binding capacity. Samples were run on a 7.5-17.5% split SDS-PAGE gel and blotted for RhoA; Monoclonal Mouse anti-RhoA antibody (Cytoskeleton inc. Cat#ARH05) was used at 1:500 dilution in TBST overnight.

### Time-lapse microscopy, immunofluorescence and image analysis

Cells were treated to a final concentration of 10nM calyculin A diluted into cell media plus 10mM HEPES at the start of time-lapse imaging. Phase contrast images for timecourse movies were taken every 4 minutes for an hour using a 20x objective on a Zeiss AXIO widefield inverted microscope fitted with a 37°C temperature environmental chamber. For blebbistatin experiments, blebbistatin was added to the media to a final concentration of 25uM, 30 minutes prior to addition of calyculin A. For of calyculin A treatment timecourse movies and still frames, imaging was started immediately before addition of calyculin A and time zero was defined as the first captured frame after calyculin A addition.

Prior to fixation for immunofluorescence, Jeg3 cells grown on coverslips were washed with PBS and pre-stained with WGA at 1:300 for 30 minutes. For actin turnover experiments in Figure 5A, Jeg3 cells were treated with either DMSO, 500nM Jasplakinolide or 100ng/mL Latrunculin B for 30 min, prior to fixation. Otherwise, for all other immunofluorescent experiments, cells grown on glass coverslips were fixed in 3.7% formalin/PBS for 15 min at room temperature. Cells were then washed with PBS and blocked with immunofluorescence buffer (PBS + 0.5% BSA + 0.5% goat serum + 0.1% Triton X-100) for 10 min. Primary and secondary antibodies were then applied in immunofluorescence buffer containing 2% FBS. Alexa Fluor–conjugated phalloidin, was added to the secondary. The cells were mounted in SlowFade Diamond Antifade (Thermo Fisher) and imaged using a spinning-disk (Yokogawa CSU-X1; Intelligent Imaging Innovations) Leica DMi600B microscope with spherical aberration correction device and a 100/1.46 numerical aperture Leica objective. Images were acquired with a metal-oxide semiconductor device camera (sCMOS) and image slices were assembled using SlideBook 6 software (Intelligent Imaging Innovations). Maximum or summed intensity projections were assembled in SlideBook 6 and exported to Illustrator software (Adobe). For clarity, side projections were vertically expanded using Illustrator. Deconvolution images were captured using an inverted Leica DMi8 wide-field microscope equipped a Leica 506249 100X/1.49 oil objective, Leica DFC 9000 GTC camera, with Leica application suit X thunder deconvolution software.

The presence or absence of microvilli was scored as described previously (Garbett et al., 2010; Hanono et al., 2006; LaLonde et al., 2010; Pelaseyed et al., 2017; Sauvanet et al., 2015). More than 50 cells per replicate were stained using the WGA, ezrin and phalloidin and binned into two categories: microvilli and no microvilli. Microvilli above cell–cell junctions were ignored in the scoring.

For calculating tortuosity among tight junctions and relative apical and lateral actin intensities, cells were stained with a ZO-1 antibody and phalloidin and imaged on a spinning–disk microscope as described earlier. To analyze tortuosity, the ZO-1 channel was skeletonized into 1 pixel line widths using ImageJ. Between two consecutive intersection points, a straight-line length was measured and the average of two manually traced line lengths (to account for human variability) along the skeletonized line were also measured. From there, the ratio between the actual line length to the expected line length (straight length) was calculated and exported into Graphpad/Prism for statistical analysis. For comparing actin intensities between apical and lateral regions, Max-Z projections were divided between ZO-1 positive (apical) and ZO-1 negative (lateral) staining. From there, outlines of each cell were traced and measured for their relative mean intensity value in the apical stacks versus the lateral stacks. The ratio between the apical and lateral values were then calculated, plotted and analyzed for statistical significance (Welch’s t-test) in GraphPad/Prism.

### Atomic Force Microscopy (AFM)

Atomic force microscopy experiments were performed using a MFP-3D-BIOTM Atomic Force Microscope by Asylum Research mounted on an Olympus IX71 inverted microscope that resides on a Herzan AVI 350-S Active Vibration Isolation Table totally enclosed in an air tight BCH-45 Large Acoustic Isolation Hood. Software modules are written in Igor Pro by Wavemetrics. Cells were plated onto WillCo-Dish Glass bottom dish (size 50×7mm. glass 30mm. Class #1.5, 0.17mm (Product number GWST-5030; Willco Wells)). Pyramidal shaped Probe TIP (PNP-TR-20) from Asylum Research (804.NW.PNP-TR) were used and calibrated using an in-air calculation of the spring constant. Individual cells within monolayers were selected using phase contrast live imaging with the probe tip location simultaneously illuminated onto the field of view. Probe tips were lowered onto a cell at a velocity of 1000nm/s to a trigger point of 1 nN. Individual cells were force mapped by indenting 100 times over a square 20μm x 20μm area. Simultaneous force indentation and membrane retraction measurements were made by first incubating probe tips with 3.0mg/ml of concanavalin A in PBS buffer for 1.5 hours at room temperature. Deflection and z-probe position data was exported into matlab then analyzed using the open access software from Huth and colleagues (Huth et al., 2019) modified for a pyramidal shape probe tip. Force indentation traces were filtered for single force peaks then chi-value goodness fit to the Hertz model. Youngs modulus statistical analysis and graphing of AFM data was performed in Graphpad/Prism.

## Acknowledgements

We thank the University of Vermont College of Medicine Microscopy Imaging Center, specifically D. Taatjes and N. Bouffard for their help using the Asylum Research MFP-3 D BIO atomic force microscope. The microscope was purchased with support from NIH award number S10RR025498 from the National Center for Research Resources (to DT). We thank Alan Howe, Robert Bauer and Albert van der Vliet (University of Vermont) for their help with AFM data collection and suggestions on Myosin Light Chain detection and Steven Huth (Christian-Albrechts University of Keil) for his help implementing AFM Hertz fit analysis to our data. We thank Jan Lammerding (Cornell University) for his generosity with reagents and use of equipment and Adam Martin (MIT) for his helpful discussions and reviewing the manuscript. We appreciate the effort by Shannon Marshall and Marcus Smolka (Cornell University) with attempts to identify additional LOK phosphorylation targets. A.T.L. was supported by a Fleming Research Fellowship. This work was supported by NIH Grants RO1 GM036552 and R35 GM131751 to A.B.

## SUPPLEMENTAL MATERIALS

**Figure S1:**
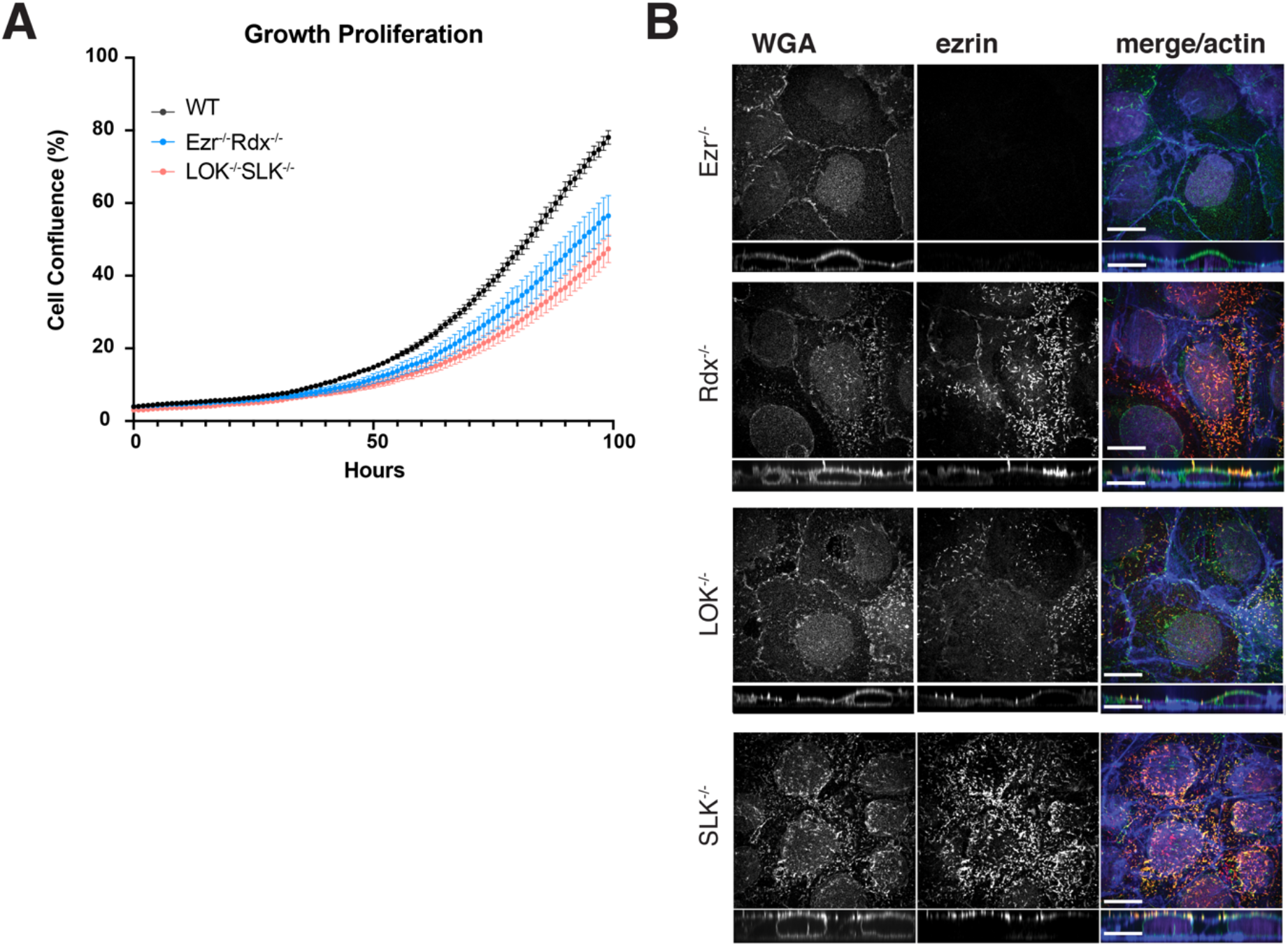
Single KO phenotypes in Jeg3 cells. (A) Growth curve WT (grey) vs. Ezr^-/-^Rad^-/-^ (Blue) and LOK^-/-^SLK^-/-^ cells (orange). N=15 wells of cells shown WT ; N=15 Ezr^-/-^Rad^-/-^; N=15 LOK^-/-^SLK^-/-^. 3,000 cells were seeded per well on 96-well plates and imaged every hour using Incucyte Live-cell analysis system. Each point represents cell confluence, measured by Incucyte over time per hour, from 0 to 100 hours. Error bars = SEM (B) Representative immunofluorescent staining of microvilli in single KO cells used for microvilli quantification and assessment of partial defects.

**Figure S2:**
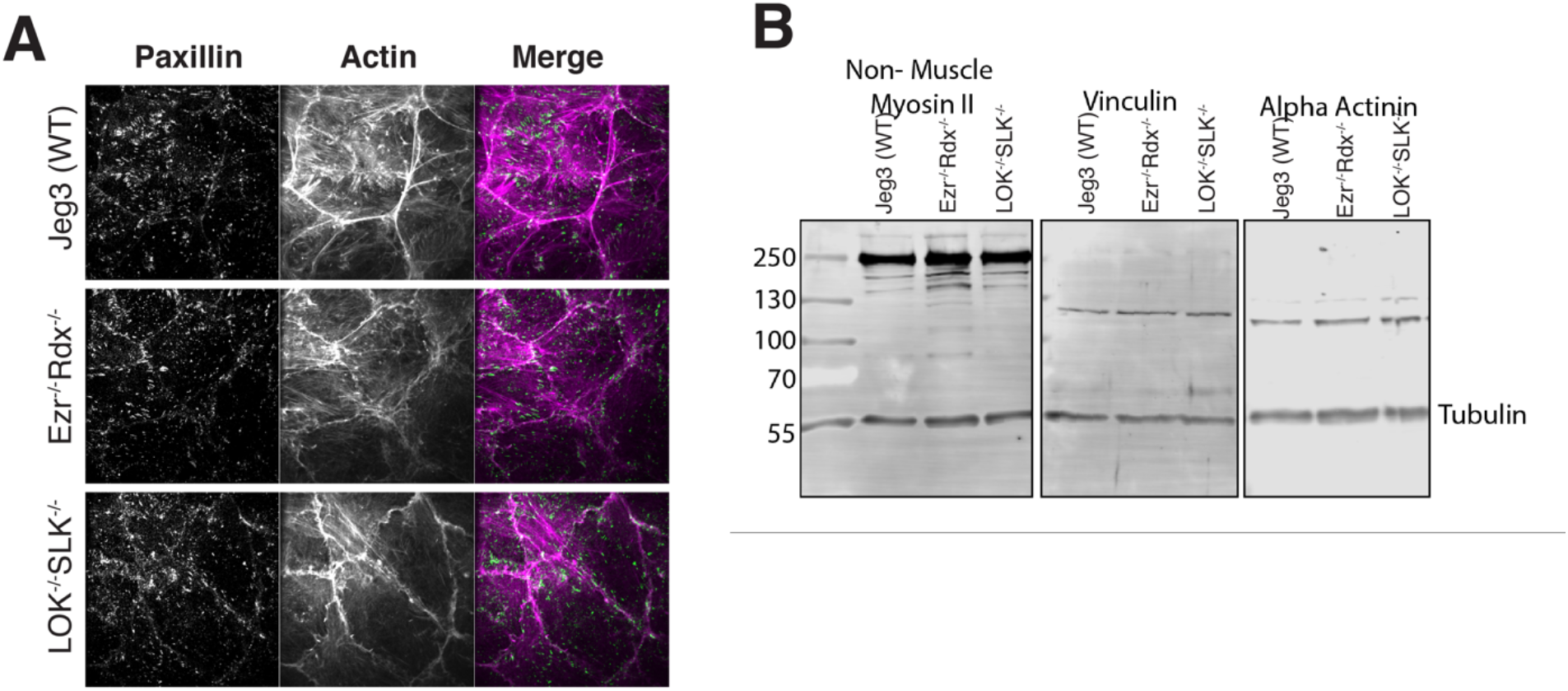
Focal contacts in WT Jeg3 vs LOK^-/-^SLK^-/-^, Ezr^-/-^Rad^-/-^ cells. (A) Immunofluorescent staining of Paxillin focal adhesion marker and actin shows similar organization. (B) Western blotting of Non-Muscle Myosin II, Vinculin and alpha-actinin and tubulin in Jeg3 WT and double knockout cells.

**Figure S5:**
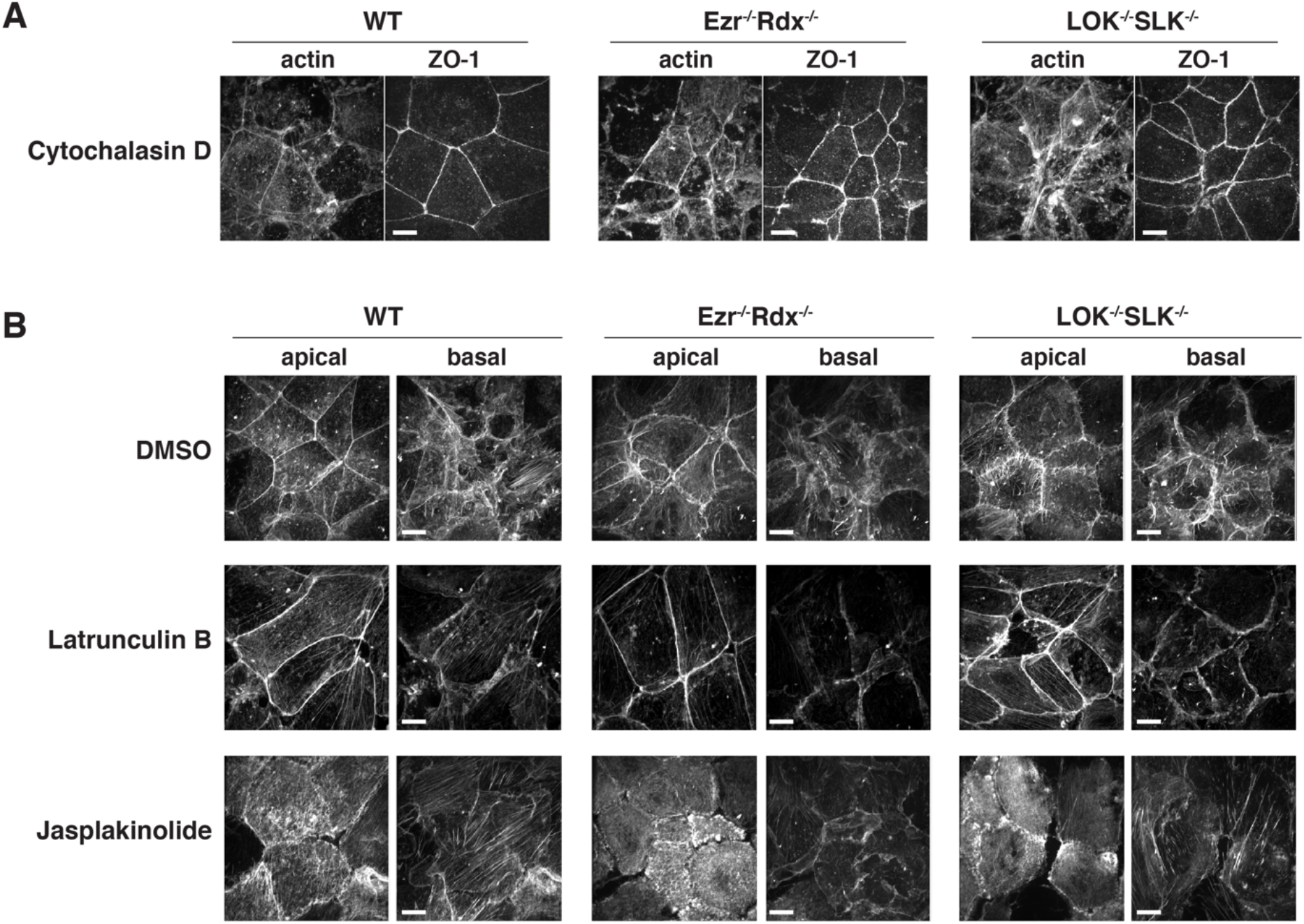
Effects of actin polymerizing drugs on Jeg-3 epithelial cells. (A) Jeg3 cells treated with 1μg/mL of Cytochalasin D for 30 min prior to fixation and staining with ZO-1 and actin. (B) Actin staining of apical and basolateral slices from maximum projection images in Figure 5A. Notably, Latrunculin B treatment favors disassembly of basolateral actin over apical actin, while maintaining a strong presence of junctional actin. Scale bar 10μm.

**Movie S1:** Time course movie of Jeg3 WT, Ezr^-/-^Rdx^-/-^ and LOK^-/-^SLK^-/-^ cells treated with 10nM calyculin A over 1 hour.

**Movie S2:** Time course movie of Jeg3 WT, and Ezr^-/-^Rdx^-/-^ LOK^-/-^SLK^-/-^ cells treated with 10nM calyculin A and 25uM blebbistatin over 1 hour. Blebbistatin treatment was added 30 minutes prior to movie start.

**Movie S3:** Time course movie of Ezr^-/-^Rdx^-/-^ and LOK^-/-^SLK^-/-^ cells transfected with phosphomimetic ezrin (Ezrin-T567E) treated with 10nM calyculin A over 1 hour.

## CITATIONS

Arber S, Barbayannis FA, Hanser H, Schnelder C, Stanyon CA, Bernards O, Caroni P. 1998. Regulation of actin dynamics through phosphorylation of cofilin by LIM-kinase. Nature 393:805–809. doi:10.1038/31729

Arnold TR, Shawky JH, Stephenson RE, Dinshaw KM, Higashi T, Huq F, Davidson LA, Miller AL. 2019. Anillin regulates epithelial cell mechanics by structuring the medial-apical actomyosin network. Elife 8:39065. doi:10.7554/eLife.39065

Bagci H, Sriskandarajah N, Robert A, Boulais J, Elkholi IE, Tran V, Lin Z-YY, Thibault M-PP, Dubé N, Faubert D, Hipfner DR, Gingras A-CC, Côté J-FF. 2020. Mapping the proximity interaction network of the Rho-family GTPases reveals signalling pathways and regulatory mechanisms. Nat Cell Biol 22:120–134. doi:10.1038/s41556-019-0438-7

Baumgartner M, Sillman AL, Blackwood EM, Srivastava J, Madson N, Schilling JW, Wright JH, Barber DL. 2006. The Nck-interacting kinase phosphorylates ERM proteins for formation of lamellipodium by growth factors. Proc Natl Acad Sci U S A 103:13391–13396. doi:10.1073/pnas.0605950103

Belkina N V., Liu Y, Hao J-JJJ, Karasuyama H, Shaw S. 2009. LOK is a major ERM kinase in resting lymphocytes and regulates cytoskeletal rearrangement through ERM phosphorylation. Proc Natl Acad Sci U S A 106:4707–4712. doi:10.1073/pnas.0805963106

Bretscher A. 1989. Rapid phosphorylation and reorganization of ezrin and spectrin accompany morphological changes induced in A-431 cells by epidermal growth factor. J Cell Biol 108:921–930. doi:10.1083/jcb.108.3.921

Bretscher A, Chambers D, Nguyen R, Reczek D. 2000. ERM-Merlin and EBP50 Protein Families in Plasma Membrane Organization and Function. Annu Rev Cell Dev Biol 16:113–143. doi:10.1146/annurev.cellbio.16.1.113

Chiasson-Mackenzie C, Morris ZS, Liu CH, Bradford WB, Koorman T, McClatchey AI. 2018. Merlin/ERM proteins regulate growth factor-induced macropinocytosis and receptor recycling by organizing the plasma membrane:Cytoskeleton interface. Genes Dev 32:1201–1214. doi:10.1101/gad.317354.118

Clucas J, Valderrama F. 2015. ERM proteins in cancer progression. J Cell Sci 128:1253–1253. doi:10.1242/jcs.170027

Fehon RG, McClatchey AI, Bretscher A. 2010. Organizing the cell cortex: The role of ERM proteins. Nat Rev Mol Cell Biol 11:276–287. doi:10.1038/nrm2866

Fukata Y, Kimura K, Oshiro N, Saya H, Matsuura Y, Kaibuchi K. 1998. Association of the myosin-binding subunit of myosin phosphatase and moesin: Dual regulation of moesin phosphorylation by Rho-associated kinase and myosin phosphatase. J Cell Biol 141:409–418. doi:10.1083/jcb.141.2.409

Garbett D, LaLonde DP, Bretscher A. 2010. The scaffolding protein EBP50 regulates microvillar assembly in a phosphorylation-dependent manner. J Cell Biol 191:397–413. doi:10.1083/jcb.201004115

Gary R, Bretscher A. 1995. Ezrin self-association involves binding of an N-terminal domain to a normally masked C-terminal domain that includes the F-actin binding site. Mol Biol Cell 6:1061–75. doi:10.1091/mbc.E12-02-0152

Gloerich M, Ten Klooster JP, Vliem MJ, Koorman T, Zwartkruis FJ, Clevers H, Bos JL. 2012. Rap2A links intestinal cell polarity to brush border formation. Nat Cell Biol 14:793–801. doi:10.1038/ncb2537

Haas MA, Vickers JC, Dickson TC. 2007. Rho kinase activates ezrin-radixin-moesin (ERM) proteins and mediates their function in cortical neuron growth, morphology and motility in vitro. J Neurosci Res 85:34–46. doi:10.1002/jnr.21102

Hall A. 1998. Rho GTpases and the actin cytoskeleton. Science (80-) 279:509–514. doi:10.1126/science.279.5350.509

Hall A, Nobes CD. 2000. Rho GTPases: Molecular switches that control the organization and dynamics of the actin cytoskeleton. Philos Trans R Soc B Biol Sci 355:965–970. doi:10.1098/rstb.2000.0632

Hanono A, Garbett D, Reczek D, Chambers DN, Bretscher A. 2006. EPI64 regulates microvillar subdomains and structure. J Cell Biol 175:803–813. doi:10.1083/jcb.200604046

Hartsock A, Nelson WJ. 2008. Adherens and tight junctions: Structure, function and connections to the actin cytoskeleton. Biochim Biophys Acta - Biomembr. doi:10.1016/j.bbamem.2007.07.012

Hayashi K, Yonemura S, Matsui T, Tsukita S. 1999. Immunofluorescence detection of ezrin/radixin/moesin (ERM) proteins with their carboxyl-terminal threonine phosphorylated in cultured cells and tissues. J Cell Sci 112:1149–1158.

Hebert AM, DuBoff B, Casaletto JB, Gladden AB, McClatchey AI. 2012. Merlin/ERM proteins establish cortical asymmetry and centrosome position. Genes Dev 26:2709–2723. doi:10.1101/gad.194027.112

Hipfner DR, Cohen SM. 2003. The Drosophila sterile-20 kinase slik controls cell proliferation and apoptosis during imaginal disc development. PLoS Biol 1:244–256. doi:10.1371/journal.pbio.0000035

Hipfner DR, Keller N, Cohen SM. 2004. Slik Sterile-20 kinase regulates Moesin activity to promote epithelial integrity during tissue growth. Genes Dev 18:2243–2248. doi:10.1101/gad.303304

Ishihara H, Martin BL, Brautigan DL, Karaki H, Ozaki H, Kato Y, Fusetani N, Watabe S, Hashimoto K, Uemura D, Hartshorne DJ. 1989. Calyculin A and okadaic acid: Inhibitors of protein phosphatase activity. Biochem Biophys Res Commun 159:871–877. doi:10.1016/0006-291X(89)92189-X

Jankovics F, Sinka R, Lukácsovich T, Erdélyi M. 2002. MOESIN crosslinks actin and cell membrane in Drosophila oocytes and is required for OSKAR anchoring. Curr Biol 12:2060–2065. doi:10.1016/S0960-9822(02)01256-3

Kimura K, Ito M, Amano M, Chihara K, Fukata Y, Nakafuku M, Yamamori B, Feng J, Nakano T, Okawa K, Iwamatsu A, Kaibuchi K. 1996. Phosphorylation and activation of myosin by Rho-associated kinase (Rho-kinase). J Biol Chem 271:20246–20249. doi:10.1074/jbc.271.34.20246

Kuramochi S, Moriguchi T, Kuida K, Endo J, Semba K, Nishida E, Karasuyama H. 1997. LOK is a novel mouse STE20-like protein kinase that is expressed predominantly in lymphocytes. J Biol Chem 272:22679–22684. doi:10.1074/jbc.272.36.22679

LaLonde DP, Garbett D, Bretscher A. 2010. A regulated complex of the scaffolding proteins PDZK1 and EBP50 with ezrin contribute to microvillar organization. Mol Biol Cell 21:1519–29. doi:10.1091/mbc.E10-01-0008

Loomis PA, Zheng L, Sekerková G, Changyaleket B, Mugnaini E, Bartles JR. 2003. Espin cross-links cause the elongation of microvillus-type parallel actin bundles in vivo. J Cell Biol 163:1045–1055. doi:10.1083/jcb.200309093

Maekawa H, Neuner A, Rüthnick D, Schiebel E, Pereira G, Kaneko Y. 2017. Polo-like kinase Cdc5 regulates Spc72 recruitment to spindle pole body in the methylotrophic yeast ogataea polymorpha. Elife 6:1–28. doi:10.7554/eLife.24340

Marjoram RJ, Lessey EC, Burridge K. 2014. Regulation of RhoA Activity by Adhesion Molecules and Mechanotransduction. Curr Mol Med 14:199–208. doi:10.2174/1566524014666140128104541

Matsui T, Maeda M, Doi Y, Yonemura S, Amano M, Kaibuchi K, Tsukita Sachiko, Tsukita Shoichiro. 1998. Rho-kinase phosphorylates COOH-terminal threonines of ezrin/radixin/moesin (ERM) proteins and regulates their head-to-tail association. J Cell Biol 140:647–657. doi:10.1083/jcb.140.3.647

Matsui T, Yonemura S, Tsukita SS, Tsukita SS. 1999. Activation of ERM proteins in vivo by Rho involves phosphatidylinositol 4-phosphate 5-kinase and not ROCK kinases. Curr Biol 9:1259–1262. doi:10.1016/s0960-9822(99)80508-9

McCormack J, Welsh NJ, Braga VMM. 2013. Cycling around cell-cell adhesion with rho GTPase regulators. J Cell Sci. doi:10.1242/jcs.097923

Meenderink LM, Gaeta IM, Postema MM, Cencer CS, Chinowsky CR, Krystofiak ES, Millis BA, Tyska MJ. 2019. Actin Dynamics Drive Microvillar Motility and Clustering during Brush Border Assembly. Dev Cell 50:545–556.e4. doi:10.1016/j.devcel.2019.07.008

Neisch AL, Formstecher E, Fehon RG. 2013. Conundrum, an ARHGAP18 orthologue, regulates RhoA and proliferation through interactions with Moesin. Mol Biol Cell 24:1420–1433. doi:10.1091/mbc.E12-11-0800

Ng T, Parsons M, Hughes WE, Monypenny J, Zicha D, Gautreau A, Arpin M, Gschmeissner S, Verveer PJ, Bastiaens PIH, Parker PJ. 2001. Ezrin is a downstream effector of trafficking PKC-integrin complexes involved in the control of cell motility. EMBO J 20:2723–2741. doi:10.1093/emboj/20.11.2723

Oshiro N, Fukata Y, Kaibuchi K. 1998. Phosphorylation of moesin by Rho-associated kinase (Rho-kinase) plays a crucial role in the formation of microvilli-like structures. J Biol Chem 273:34663–34666. doi:10.1074/jbc.273.52.34663

Pakkanen R, Hedman K, Turunen O. 1987. Microvillus-specific M 75,000 plasma membrane protein of human choriocarcinoma cells. J Histochem Cytochem 35:809–816. doi:10.1177/35.8.3298422

Pearson MA, Reczek D, Bretscher A, Karplus PA. 2000. Structure of the ERM protein moesin reveals the FERM domain fold masked by an extended actin binding tail domain. Cell 101:259–270. doi:10.1016/S0092-8674(00)80836-3

Pelaseyed T, Sauvanet C, Viswanatha R, Filter JJ, Goldberg ML, Bretscher A. 2017. Ezrin activation by LOK phosphorylation involves a PIP2-dependent wedge mechanism. Elife 6:1–18. doi:10.7554/eLife.22759

Pietromonaco SF, Simons PC, Altman A, Elias L. 1998. Protein kinase C-θ phosphorylation of moesin in the actin-binding sequence. J Biol Chem 273:7594–7603. doi:10.1074/jbc.273.13.7594

Ratheesh A, Gomez GA, Priya R, Verma S, Kovacs EM, Jiang K, Brown NH, Akhmanova A, Stehbens SJ, Yap AS. 2012. Centralspindlin and alpha-catenin regulate Rho signalling at the epithelial zonula adherens. Nat Cell Biol 14:818–828. doi:10.1038/ncb2532

Reczek D, Berryman M, Bretscher A. 1997. Identification of EPB50: A PDZ-containing phosphoprotein that associates with members of the ezrin-radixin-moesin family. J Cell Biol 139:169–179. doi:10.1083/jcb.139.1.169

Reyes CC, Jin M, Breznau EB, Espino R, Delgado-Gonzalo R, Goryachev AB, Miller AL. 2014. Anillin regulates cell-cell junction integrity by organizing junctional accumulation of Rho-GTP and actomyosin. Curr Biol 24:1263–1270. doi:10.1016/j.cub.2014.04.021

Rodriguez-Boulan E, Macara IG. 2014. Organization and execution of the epithelial polarity programme. Nat Rev Mol Cell Biol 15:225–242. doi:10.1038/nrm3775

Sanjana NE, Shalem O, Zhang F. 2014. Improved vectors and genome-wide libraries for CRISPR screening. Nat Methods. doi:10.1038/nmeth.3047

Sanson M, Marineau C, Desmaze C, Lutchman M, Ruttledge M, Baron C, Narod S, Delattre O, Lenoir G, Thomas G, Aurlas A, Rouleau GA. 1993. Germline deletion in a neurofibromatosis type 2 kindred inactivates the NF2 gene and a candidate meningioma locus. Hum Mol Genet 2:1215–1220. doi:10.1093/hmg/2.8.1215

Saotome I, Curto M, McClatchey AI. 2004. Ezrin is essential for epithelial organization and villus morphogenesis in the developing intestine. Dev Cell 6:855–864. doi:10.1016/j.devcel.2004.05.007

Sauvanet C, Garbett D, Bretscher A. 2015. The function and dynamics of the apical scaffolding protein E3KARP are regulated by cell-cycle phosphorylation. Mol Biol Cell 26:3615–3627. doi:10.1091/mbc.E15-07-0498

Shalem O, Sanjana NE, Hartenian E, Shi X, Scott DA, Mikkelsen TS, Heckl D, Ebert BL, Root DE, Doench JG, Zhang F. 2014. Genome-scale CRISPR-Cas9 knockout screening in human cells. Science (80-) 343:84–87. doi:10.1126/science.1247005

Speck O, Hughes SC, Noren NK, Kulikauskas RM, Fehon RG. 2003. Moesin functions antagonistically to the Rho pathway to maintain epithelial integrity. Nature 421:83–87. doi:10.1038/nature01295

ten Klooster JP, Jansen M, Yuan J, Oorschot V, Begthel H, Di Giacomo V, Colland F, de Koning J, Maurice MM, Hornbeck P, Clevers H. 2009. Mst4 and Ezrin Induce Brush Borders Downstream of the Lkb1/Strad/Mo25 Polarization Complex. Dev Cell 16:551–562. doi:10.1016/j.devcel.2009.01.016

Terry SJ, Zihni C, Elbediwy A, Vitiello E, San IVLC, Balda MS, Matter K. 2011. Spatially restricted activation of RhoA signalling at epithelial junctions by p114RhoGEF drives junction formation and morphogenesis. Nat Cell Biol 13:159–166. doi:10.1038/ncb2156

Tran Quang C. 2000. Ezrin function is required for ROCK-mediated fibroblast transformation by the Net and Dbl oncogenes. EMBO J 19:4565–4576. doi:10.1093/emboj/19.17.4565

Trofatter JA, MacCollin MM, Rutter JL, Murrell JR, Duyao MP, Parry DM, Eldridge R, Kley N, Menon AG, Pulaski K, Haase VH, Ambrose CM, Munroe D, Bove C, Haines JL, Martuza RL, MacDonald ME, Seizinger BR, Short MP, Buckler AJ, Gusella JF. 1993. A novel moesin-, ezrin-, radixin-like gene is a candidate for the neurofibromatosis 2 tumor suppressor. Cell 72:791–800. doi:10.1016/0092-8674(93)90406-G

Turunen O, Wahlström T, Vaheri A. 1994. Ezrin has a COOH-terminal actin-binding site that is conserved in the ezrin protein family. J Cell Biol 126:1445–1453. doi:10.1083/jcb.126.6.1445

Velasco G, Armstrong C, Morrice N, Frame S, Cohen P. 2002. Phosphorylation of the regulatory subunit of smooth muscle protein phosphatase 1M at Thr850 induces its dissociation from myosin. FEBS Lett 527:101–104. doi:10.1016/S0014-5793(02)03175-7

Viswanatha R, Ohouo PY, Smolka MB, Bretscher A. 2012. Local phosphocycling mediated by LOK/SLK restricts ezrin function to the apical aspect of epithelial cells. J Cell Biol 199:969–984. doi:10.1083/jcb.201207047

Yang N, Higuchi O, Ohashi K, Nagata K, Wada A, Kangawa K, Nishida E, Mizuno K. 1998. Cofilin phosphorylation by LIM-kinase 1 and its role in Rac-mediated actin reorganization. Nature 393:809–812. doi:10.1038/31735

Zihni C, Balda MS, Matter K. 2014. Signalling at tight junctions during epithelial differentiation and microbial pathogenesis. J Cell Sci 127:3401–3413. doi:10.1242/jcs.145029

